# Biomolecular condensates of Eya drive transcriptional co-activation associated with eye development

**DOI:** 10.1101/2025.10.15.682020

**Authors:** Nilanjana Das, Janhavi Bapat, Amitabha Majumdar

## Abstract

Eyes absent (Eya), a transcriptional coactivator essential for eye development in Drosophila, also plays an important role in organ development in mammals and is associated with several diseases. To better understand the mechanism of Eya-mediated transcriptional co-activation, we find that Eya forms biomolecular condensates with liquid-like properties. These condensates function as possible transcription hubs, through the compartmentalization of key eye development regulators, the transcription factors So, Dac, Optix, Ey, and the kinase Nemo, RNA Pol II machinery, P300/CBP, and show the presence of target RNAs. We map the PST-TPM-PST region of Eya, which is critical for transcriptional co-activation and eye development, to be crucial for condensate formation as well. We identify a human deafness-associated mutation mapping to a conserved Drosophila site in the PST-TPM-PST region, which impairs transcriptional co-activation and shifts the material property of the Eya condensates to a less dynamic state. Our findings thus provide evidence that the condensation of Eya and its dynamic state are integral to transcription regulation and its role in development and disease mechanisms.

## Introduction

The nuclear environment is densely packed with a lot of nuclear proteins, RNA, and DNA, along with membrane-less organelles like nucleolus, Cajal bodies, nuclear speckles, etc. For precise nuclear function, all these need to be spatially organized while maintaining their boundaries without mixing with their surroundings. Recent studies have suggested that many of the membrane-less compartments originate through a process of liquid-liquid phase separation (LLPS) akin to the demixing of oil and water. Post formation of these compartments, also now referred to as biomolecular condensates, they often behave like liquid droplets, creating chemically distinct microenvironments within the cell. The advantage of such condensates is that this enables the concentration of the specific biomolecules in a noncovalent and reversible manner, enabling the containment of zones for various localized biochemical processes, and yet being able to maintain active exchange with the surroundings. These dynamic condensates undergo fusion when they come close to each other and can undergo rapid internal rearrangement of molecules, due to their liquid-like nature (Shin & Brangwynne, 2017; Banani *et al*, 2017; Boeynaems *et al*, 2018). The presence of intrinsically disordered regions (IDRs) in the protein sequences and multivalent interactions amongst the proteins are suggested to be the drivers for the formation of biomolecular condensates through LLPS (Banani *et al*, 2017; Borcherds *et al*, 2021).

LLPS-mediated nuclear sub-compartmentalization has emerged as a fundamental principle in the maintenance of the nuclear organization. SPOP, a component of nuclear speckles, undergoes LLPS, and this is essential for its localization in nuclear speckles (Marzahn *et al*, 2016; Bouchard *et al*, 2018). The Nucleolus made of a mix of RNA and protein, the site for ribosome subunit biogenesis, RNP assembly, behaves like a phase-separated condensate, where nucleolar proteins phase separate to form coexisting immiscible phases that form different sub-compartments with non-overlapping functions (Boisvert *et al*, 2007; Lafontaine *et al*, 2021; Brangwynne *et al*, 2011; Feric *et al*, 2016; King *et al*, 2024). The chromatin is also organized into subdomains, which are transcriptionally silent (heterochromatin) or active (euchromatin). Reconstituted chromatin undergoes phase separation to form dense condensates in vitro with a similar nucleosome concentration as in vivo, and the chromatin can be further sub-compartmentalized through the interaction of acetylated chromatin with Brd4 (Gibson *et al*, 2019). Heterochromatin formation is suggested to be regulated through the phase separation of HP1 protein, and histone modification (Larson *et al*, 2017; Strom *et al*, 2017; Wang *et al*, 2019). Polycomb complexes involved in gene repression have been shown to form condensates in which their ligands can partition and regulate the compaction of chromatin (Brown *et al*, 2023; Tatavosian *et al*, 2019; Kim *et al*, 2023; Plys *et al*, 2019; Eeftens *et al*, 2021).

For the euchromatin regions, precise coordination and control of transcription is regulated through a multistep mechanism, and phase separation has been suggested to be important for this (Cramer, 2019). Transcription is carried out by RNA pol II but requires the participation of several other proteins, including transcription factors. Transcription factors (TFs) are proteins that bind to specific sequences/motifs in the promoter and enhancers in DNA and regulate transcription, and are thought to be the master regulators behind cell-type or tissue-specific gene expression. TFs can make the chromatin accessible by remodeling the nucleosomes, assembling the pre-initiation complex, phosphorylation of the CTD of RNA pol II, promoter clearance, and elongation, etc. Transcription in certain cases needs transcription coactivators that do not bind to DNA but regulate transcription by (a) binding to transcription factors and regulating the transcription of the target genes (b) acting as scaffolds to recruit other proteins with enzymatic activities needed to modify the chromatin accessibility by either histone modification or through ATP dependent chromatin remodeling, (c) interacting with the transcription complex proteins and RNA Pol II (Spiegelman & Heinrich, 2004). Transcription factors, coactivators, and RNA Polymerase II have been reported to assemble on specific enhancers through an LLPS-like process, with both transcription factors and coactivators have been found to co-partition in condensates and regulate transcription (Sabari *et al*, 2020; Pei *et al*, 2025).

These condensate mediated control of gene activation and repression is also expected to play a crucial role in developmental processes in an organism. Evidence towards the role of condensates in developmental processes has come from diseased conditions like hereditary synpolydactyly, hand-foot genital syndrome, cleidocranial dysplasia, and spinocerebellar ataxia type 17, which are associated with repeat expansions in transcription factors. These expansions alter their phase separation property and cause a reduction in co-partitioning the co-activators, altering the composition of the condensates and dysregulation of transcription (Sabari, 2020; Mathias *et al*, 2024; Basu *et al*, 2020).

In animal development, one of the most interesting processes is the development of the eye, which is regulated by a network of gene regulatory pathways. Most important among these are the master regulator genes, which are part of the retinal determination network. The master regulator genes under a loss-of-function situation result in the absence of the eye, and under a gain-of-function condition, when ectopically expressed in non-neuronal tissues, can convert cells to retina. This suggests these genes have the powerful capacity to capture and change the development program of other cell types to differentiate them into retinal cells. Amongst the members of the retinal determination genetic network (RDGN) in Drosophila, Eyes absent (Eya) (Bonini *et al*, 1993) is a unique member that is a transcriptional coactivator which cannot bind to DNA by itself. The other members, Eyeless (Ey), Twin of eyeless (Toy), Sine ocullis (So), Dachshund (Dac), and Optix (Opt), are all transcription factors with the ability to bind to DNA. Eya is evolutionarily conserved, with homologs in mice and humans. In mice, Eya1 knockouts die at birth and show multiple developmental abnormalities, including in ear development and hearing, craniofacial and skeletal defects, absent kidneys, thymus, and parathyroid glands (Xu *et al*, 1999). Mice Eya4 knockouts show defects in hearing and ear development (Depreux *et al*, 2008). In humans, Eya homologs hEya1-4 are linked to a range of developmental disorders and diseases, such as branchiootorenal syndrome, otofaciocervical syndrome, cardiofacial syndrome, late-onset deafness, cardiomyopathy, and various cancers (Tadjuidje & Hegde, 2012; Soni *et al*, 2021).

Given its broad developmental roles and being the only exceptional member of the RDGN pathway that cannot bind DNA, we sought to understand how Eya functions in transcription regulation as a coactivator. We observed Eya to form dynamic liquid-like condensates in the nucleus, and these can recruit and partition other transcription factors, Sine ocullis, Dac, Eyeless, and Optix, along with a kinase Nemo. These all play a crucial role in eye development. The condensates are possible transcription zones, as we noted the presence of RNA PolII, CBP/P300, an acetylated histone marker for the presence of enhancers, and RNA enriched in these structures. Further, we find the PST-TPM-PST region in Eya to be necessary for condensate formation. As this same region is needed for transcription co-activation and eye development, this connects the Eya condensate formation with transcription regulation and eye development. Importantly, we examine a human deafness-associated mutation in hEya1 located in a site conserved in the Drosophila PTP region. This mutation alters the material properties of the Eya condensates from liquid-like to a less dynamic, more solid-like state, with impaired transcription activation without disrupting TF compartmentalization. Together, our findings illustrate the functional importance of transcription co-activator condensation in normal development and link its material properties to disease mechanisms.

## Results

### Eya forms liquid-like nuclear condensates

Using IUPred3, we found the presence of disordered regions in Eya. The 760 amino acid open reading frame (ORF) of the protein contains a PST (Proline-Serine-Threonine) rich transactivation domain embedded with a threonine phosphatase motif (TPM) and a C-terminal Eya (ED) domain (Figure 1A). We noticed the presence of two regions with high disorder scores, one in the N-terminal and the other towards the middle of the protein’s ORF within the PST-TPM-PST (PTP) region (Figure 1B). As disordered domains were shown to promote condensate formation, and MolPhase, a machine learning predictor (Liang *et al*, 2024), provided a very high score of 0.9994, predicting a high chance of the Eya protein to phase separate and form biomolecular condensates, we next investigated whether this protein can form condensates in cells. We tagged the Eya ORF with Venus and transfected it into Drosophila S2 cells. S2 cells do not express Eya protein, thus acting as a null host for exogenous expression. In these cells, we observed Eya to form multiple condensates. On immunostaining the cells with an antibody against Lamin, which marks the nuclear envelope, we found these condensates to be localized exclusively within the nucleus (Figure 1C). On live confocal imaging to visualize the behavior of these condensates over time, we noticed that many of these condensates undergo fusion (Figure 1D and Supplemental Video 1). We next performed Fluorescence recovery after photobleaching (FRAP) experiments to see if molecules within these condensates undergo exchange with surrounding molecules. After photobleaching, we noticed rapid recovery of the bleached region (Figure 1E, F, and Supplemental Video 2). On quantitation, we observed these condensates to have ∼60% of mobile fraction. The fusion of the condensates and their rapid recovery on photobleaching indicate these condensates have properties similar to liquid droplets.

**Figure 1.**
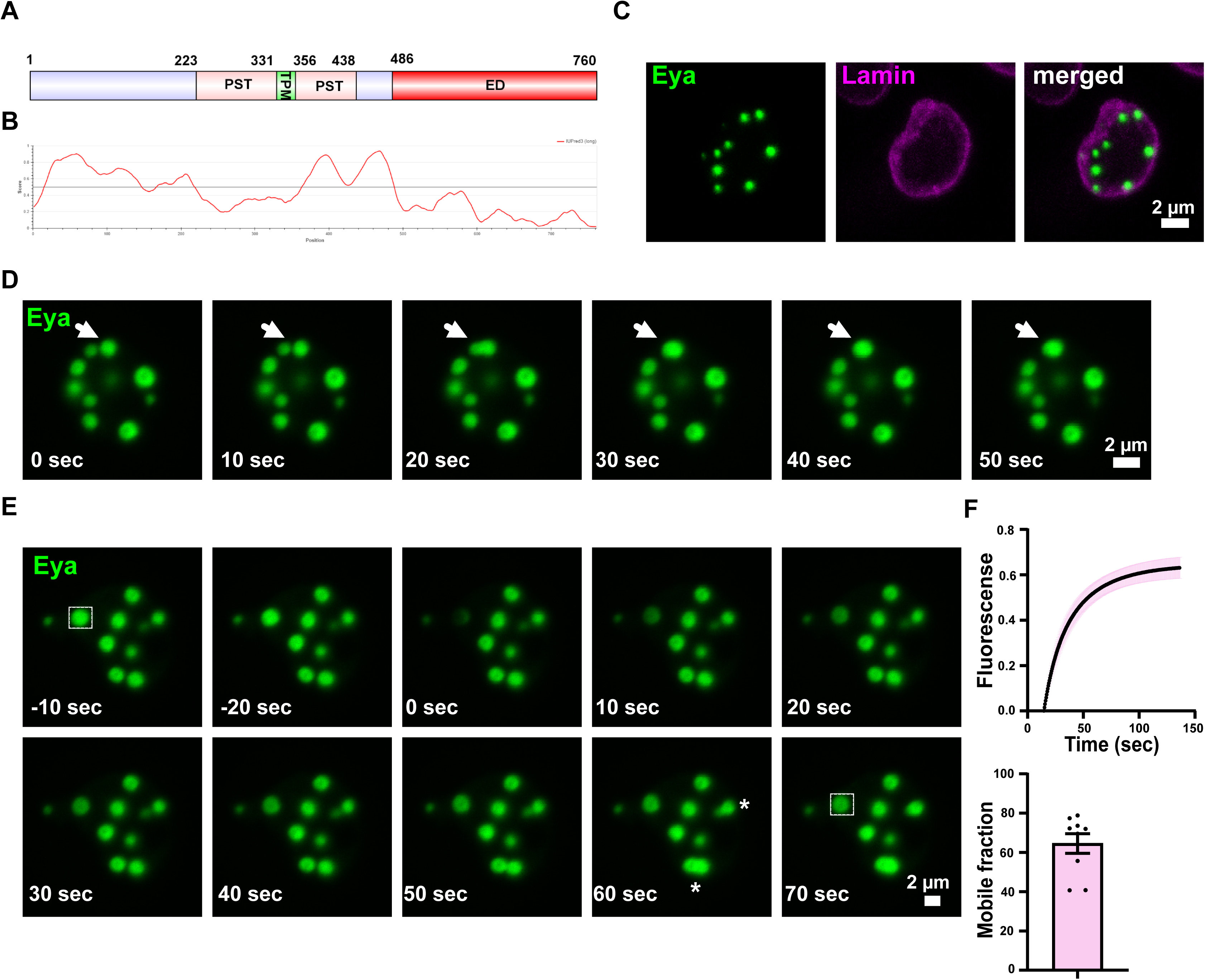
Eya forms liquid-like condensates. **A.** Schematic of the domain-based organization of Drosophila Eya **B.** IUPRED3 predicted disordered regions in Drosophila Eya **C.** Representative image of S2 cell expressing Drosophila Eya immunostained using Lamin antibody, showing condensate-like structures formed by Eya inside the nucleus **D.** Frames from live confocal imaging of S2 cell expressing Eya, showing fusion of the condensates (marked by arrow) **E.** Frames from the FRAP experiment with Eya condensates show rapid recovery of the fluorescence after photobleaching (marked by a white box at the beginning and the end). The frames also show two condensates fusing (marked by *). **F.** The top panel shows the FRAP recovery graph of Eya condensate, and the bottom panel shows the quantitation of the mobile fraction. The sample size is n=8, and the error bar represents the standard error mean (SEM).

We also found that driving the Venus-tagged Eya can rescue the eyes absent phenotype in eya^2^ (Figure 2A). This suggests that the tagging with Venus is not causing a loss of function in the Eyes absent gene.

**Figure 2.**
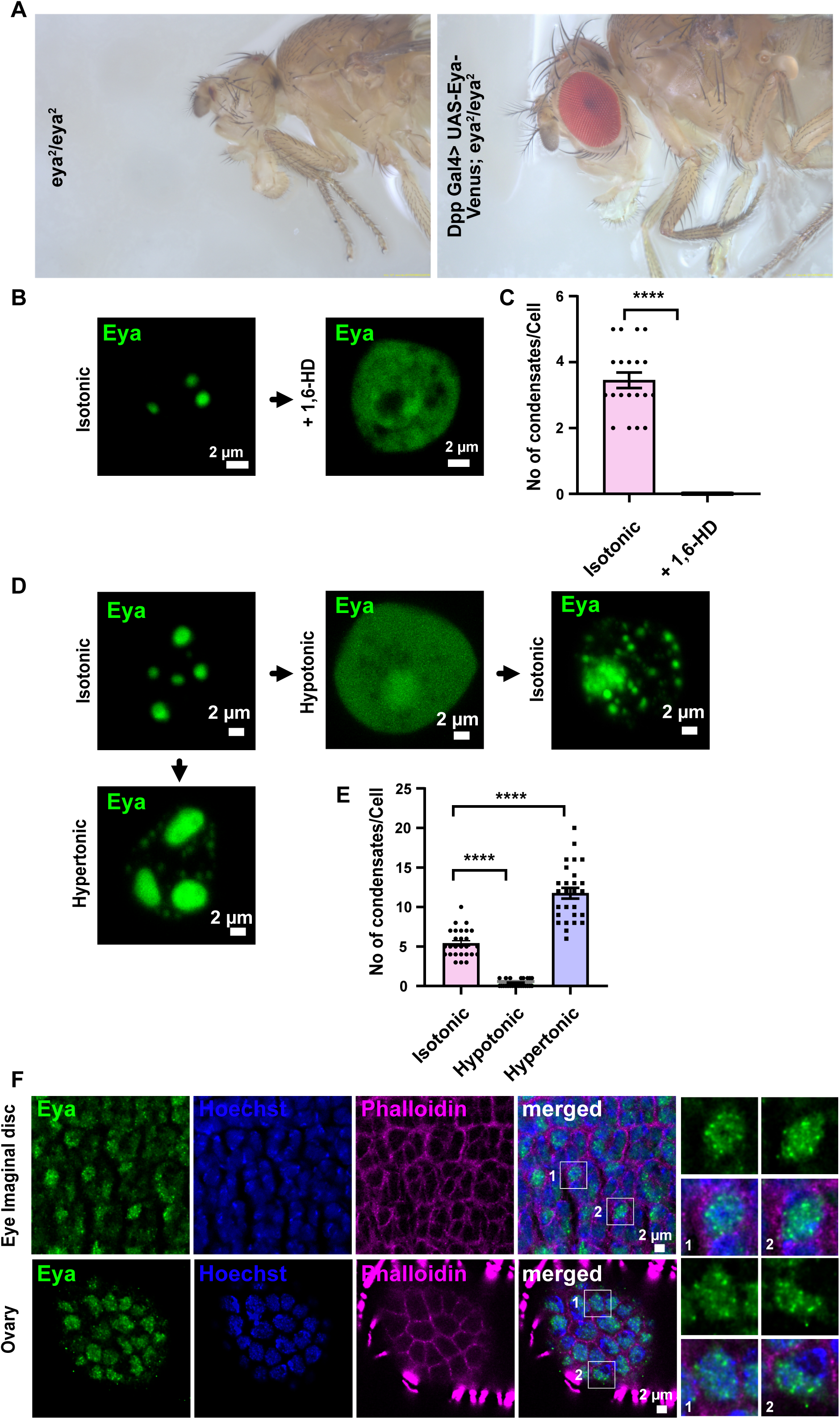
Eya condensates are responsive to osmotic challenge and 1.6-HD treatment. **A.** The eyes absent phenotype of the eya^2^ mutant can be rescued by the condensate-forming Eya-Venus construct when expressed using Dpp Gal4. **B.** Representative image of S2 cell expressing Eya in the nucleus as a control (left panel) with condensates inside the nucleus. On treatment with 1,6-Hexanediol (right panel), the condensates dissolve and fluorescence is seen both within and outside the nucleus. **C.** Quantitation of condensate number per cell showed a significant decrease on 1,6-Hexanediol treatment. Unpaired Student’s t-test was performed, where sample size (n) for the isotonic control set = 20 and n for the 1,6-HD treated set = 19. The p-value was <0.0001. Error bars represent SEMs. **D.** Representative image of S2 cell expressing Eya condensates (top left panel), which on hypotonic buffer treatment dissolves immediately (top middle panel), and on replacing this with isotonic buffer, the condensates reappear, but this time in both nucleus and cytoplasm of the cells (top right panel). On treatment with hypertonic buffer, bigger condensates are observed along with the appearance of many new smaller condensates (Bottom panel). **E.** Quantitation of condensate number per cell showing a significant decrease and increase on hypotonic and hypertonic treatment. Unpaired Student’s t-test was performed separately between Hypotonic and Hypertonic treatments to compare the differences from the isotonic control. n for isotonic, hypotonic, and hypertonic were 27, 24, and 27, respectively. P-values for both hypotonic and hypertonic were <0.0001. Error bars represent SEMs. **F.** Endogenous Eya visualized using a CRISPR-based GFP knock-in shows condensate-like structures in the cells of the eye imaginal disc and ovary.

### Eya condensates respond to 1,6-Hexanediol and osmotic perturbations

Several condensates with liquid-like nature are sensitive to treatment with an aliphatic alcohol 1,6-Hexanediol (1,6-HD) (Kroschwald *et al*, 2015; Molliex *et al*, 2015; Zheng *et al*, 2025). Hexanediol acts by disrupting the hydrophobic interactions within the condensate-forming proteins. We treated Eya expressing cells with 1,6-HD and observed the condensates to dissolve and convert to a diffused state (Figure 2B, C), supporting these condensates to be of a liquid-like nature.

Osmotic challenges have been shown to act as switches for modulating the condensate behavior of several proteins (Davis *et al*, 2019; Sehgal *et al*, 2022). We asked if Eya condensates are responsive to osmotic changes. We first subjected Eya-expressing cells to hypo-osmotic treatment. We observed that the hypo-osmotic treatment immediately dissolved the condensates to a diffused state (Figure 2D, E), and post-treatment Eya was present in both the nucleus and cytoplasm (Figure 2D). On changing the hypo-osmotic condition to a normal osmotic condition, the diffused state reverses back to the condensate state, however, now the condensates are seen in both the nucleus and cytoplasm. We also exposed these cells to hyper-osmotic treatment, and here we observed the appearance of several new, smaller condensates in the cells, suggesting this is inducing the formation of new condensates (Figure 2D, E). Overall, this suggests osmotic conditions can be a reversible switch to alter the Eya condensates in a concentration-dependent manner inside cells, as the hypo-osmotic treatment will cause a decrease in Eya concentration inside the cell by bringing in more water, and the hyperosmotic treatment will increase the Eya concentration by removing water from the inside of the cell.

### Endogenous Eya forms condensate-like structures

Previously, endogenous Eya was reported to be expressed in larval eye imaginal discs and adult female ovaries (Bai & Montell, 2002; Chang *et al*, 2013; Dai *et al*, 2017; Bonini *et al*, 1993, 1998). To examine if Drosophila Eya at endogenous levels can form condensates in vivo, we generated a GFP knock-in at the Eya locus. These flies have Eya tagged with GFP in its C-terminal in its genomic locus. We dissected the larval eye imaginal disc and ovary from the adult animals, immunostained these tissues with anti-GFP antibody, and imaged them using confocal microscopy. Here, we were able to detect small punctate structures resembling condensates in both the imaginal disc and ovary (Figure 2F), suggesting endogenous Eya can form condensates.

### The Eya condensates partition proteins involved in the eye development pathway

We next investigated if the Eya condensates are functionally relevant, and in this context, we examined if these condensates can influence the spatial organization of the Eya interacting proteins. Amongst the interactors mentioned in flybase portal (Supplementary Figure 1), we chose Abdominal A (AbdA), Abdominal B (AbdB), IKK-β, Hop, Relish (Rel), Tim, Socs44A, Sine oculis (So), Dachshund (Dac), Optix and Eyeless (Ey) and Nemo for our experiments and tagged them with CFP (Hoi *et al*, 2016; Baëza *et al*, 2015; Liu *et al*, 2012; Kenyon *et al*, 2005a; Anderson *et al*, 2012; Abrieux *et al*, 2020; Morillo *et al*, 2012).

On co-expression with Eya-Venus, we noticed no partitioning of AbdA, AbdB, IKK-β, Hop, Rel, Tim, and Socs44A in the Eya condensates (Supplementary Figure 2A, B, C, D, E, F, G, H, I, J, K, L, M, N). In contrast, So, Dac, Optix, Ey, and Nemo co-partition and colocalize in the nuclear Eya condensates (Figure 3A, B, C, D, E, F, G, H, I, J). Amongst the proteins co-partitioning with Eya, we noticed that when expressed alone, So and Optix form single foci in the nucleus, whereas Dac and Ey show condensate-like structures mainly within the nucleus, with some in the cytoplasm. Nemo is majorly present in the nucleus but also shows small punctae in the cytoplasm. Interestingly, So, Dac, Optix, and Ey are transcription factor members of the retinal determination gene network pathway, and their mutants show a very similar phenotype to Eya mutants, that of lacking eyes (Li *et al*, 2013a; Seimiya & Gehring, 2000a, 2000b; Weasner *et al*, 2007). For three of the transcription factors, Ey, So, and Dac, immunoprecipitation assays have confirmed their interaction with Eya (Jin & Mardon, 2016; Chen *et al*, 1997; Pignoni *et al*, 1997). As Eya, a transcriptional coactivator, cannot bind to DNA directly, its partitioning of transcription factors makes the condensates functionally relevant by possibly allowing the condensates to be separate spatial zones for transcription regulation through target-specific DNA binding by the transcription factors. Nemo was previously found to be a kinase for Eya, which amplifies the Eya-So transcriptional output during retinal determination (Morillo *et al*, 2012). The partitioning of Nemo possibly brings it to the spatial zone of transcription where it needs to influence Eya and So to amplify the transcriptional output. In summary, Eya condensates selectively partition and compartmentalize proteins which play a crucial role in eye development.

**Figure 3.**
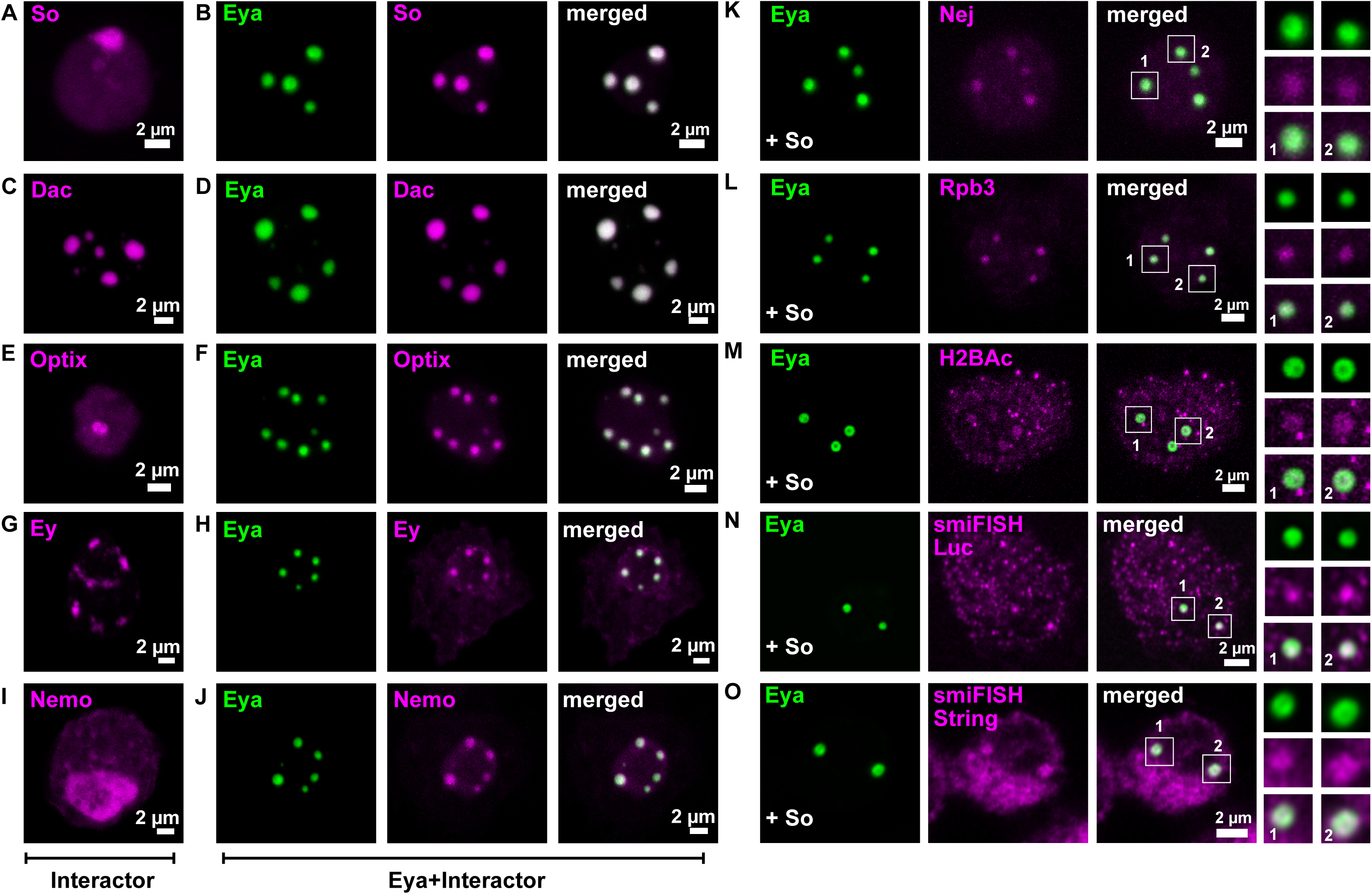
Eya condensates partition the TFs and a kinase related to eye development, and are possible transcription zones. **A-J.** Representative image of S2 cell expressing **(A)** So, **(B)** So and Eya, **(C)** Dac, **(D)** Dac and Eya, **(E)** Optix, **(F)** Optix and Eya, **(G)** Ey, **(H)** Ey and Eya, **(I)** Nemo, **(J)** Nemo and Eya. **K.** Representative image of S2 cell expressing Eya and So (not imaged) showing enrichment of endogenous Nej in the condensates. **L.** Representative image of S2 cell expressing Eya and So (not imaged) showing colocalization of RNA pol II complex’s Rpb3 subunit to colocalize in the condensates. **M.** Representative image of S2 cell expressing Eya and So (not imaged) showing enrichment of endogenous acetylated H2B in the condensates. **N.** smiFISH with probes against luciferase reporter in cells expressing Eya and So (not imaged), showing colocalization of luciferase RNA in the condensates. **O.** smiFISH with probes against luciferase reporter in cells expressing Eya and So (not imaged), showing colocalization of endogenous String RNA in the condensates.

### Eya condensates are linked to transcriptional regulation

As the Eya condensates could partition transcription factors, So, Dac, Ey, and Optix, we asked whether these condensates are involved in transcription regulation by acting as zones for transcription? We reasoned that if the Eya condensates are sites of transcriptional regulation, then the machinery associated with transcription also should be present in these condensates. As in the context of Eya mediated transcription co-activation, the most studied transcription factor is So, we focused on the Eya-So condensates.

These condensates were found not to colocalize with Histone and HP1 (Supplementary Figure 3), suggesting that these condensates are not a part of the densely packed nuclear heterochromatin regions, which are transcriptionally inactive. The Eya-So condensates also did not colocalize with Fibrillarin (Supplementary Figure 3), a marker for the nucleolus, suggesting the condensates are not part of the nucleolus.

Next, we looked at the protein CBP/P300, which plays a crucial role in transcription regulation as a co-activator through being a bridge and scaffold for the assembly of several components, including RNA PolII subunits, transcription factors, and also acting as an acetyltransferase for histones and many non-histone proteins (Chan & La Thangue, 2001; Janknecht & Hunter, 1996; Goodman & Smolik, 2000; Ogryzko *et al*, 1996; Bannister & Kouzarides, 1996). Mammalian Eya’s are earlier reported to interact with CBP/P300 (Yang & Liu, 2022; Ikeda *et al*, 2002). The CBP/P300 protein forms condensates in vivo, which co-condense with transcription factors that are directly related to the transcriptional activation of target genes and contain the newly synthesized RNAs (Ma *et al*, 2021). On immunostaining for the Drosophila homolog of P300/CBP, Nejire, we noticed the Eya-So condensates to be enriched with endogenous Nejire (Figure 3K).

We further coexpressed an RFP tagged Rpb3 construct along with Eya and So in S2 cells. Rpb3 is a subunit of the RNA polymerase II, and its RFP-tagged construct has been previously used to image transcription sites (Zobeck *et al*, 2010). On imaging, we observed Rpb3 to colocalize with the condensates (Figure 3L), suggesting the presence of the transcription machinery in these condensates. Eya condensates with Dac, Ey, and Optix also showed similar enrichment of Rpb3 and Eya condensates in the absence of transcription factors do not contain Rpb3 (unpublished data).

Transcription regulation is also controlled through the activation of enhancer elements, and acetylated H2B can be a mark of such active enhancers (Narita *et al*, 2023). On immunostaining of Eya and So coexpressing cells, we noticed the presence of acetylated H2B in the condensates (Figure 3M), indicating the association of active enhancer regions of the genome with the Eya condensates.

Next, we coexpressed Eya and So along with an 8X-ARE-Luciferase reporter construct and performed smiFISH (Tsanov *et al*, 2016) using probes against Luciferase RNA. The 8X-ARE-Luciferase reporter was previously shown to be responsive to Eya and So coexpression (Silver *et al*, 2003). We observed smiFISH signal for luciferase reporter to be present within the Eya condensates (Figure 3N). We also checked what happens to another target of Eya and So, a gene named String (Jin *et al*, 2016), which regulates cell cycle regulation and pattern formation in the retina. In smiFISH using probes against String, we noted the Eya condensates to contain the signal for endogenous String RNA (Figure 3O). Our observation with the ARE-Luciferase reporter and String RNA indicates these condensates can be sites for Eya-dependent transcriptional activation. Together, the partitioning of transcription factors, presence of Nej/P300/CBP, RNA Pol II machinery, Histone modification marker for presence of active enhancer regions, and the transcribed reporter RNA, suggest that the Eya-So condensates are associated with transcription regulation and possibly represent transcriptional hubs or zones.

### The PST-TPM-PST region of Eya is essential for condensate formation

We next aimed to map the region of Eya needed for condensate formation. We made several deletions spanning various regions and domains (Figure 4A), expressed these in S2 cells, and used microscopy to see if these constructs were forming condensates or not. Since the N-terminal of Eya had a disordered region, we made 2 deletions in this region Δ84 and Δ187. On imaging, we noticed that, similar to full-length Eya, both of these constructs form condensates (Figure 4B, C, D), suggesting the disordered region in the N-terminal is dispensable for condensate formation. Next, we imaged two deletions from the N-terminal, of 438 and 485 amino acids. In both of these Δ438 and Δ485 constructs, we noticed diffused distribution of Eya and the absence of the condensates (Figure 4E, F). In contrast, the complementary constructs containing the N-terminal 438 and 485 amino acids, Eya 438 and Eya ΔED showed the presence of condensates (Figure 4G, H). We also deleted the two internal regions corresponding to the PST-TPM-PST region (ΔPTP) and the TPM motif (ΔTPM). While the ΔTPM construct formed condensates, the ΔPTP construct showed diffused distribution (Figure 4I, J).

**Figure 4.**
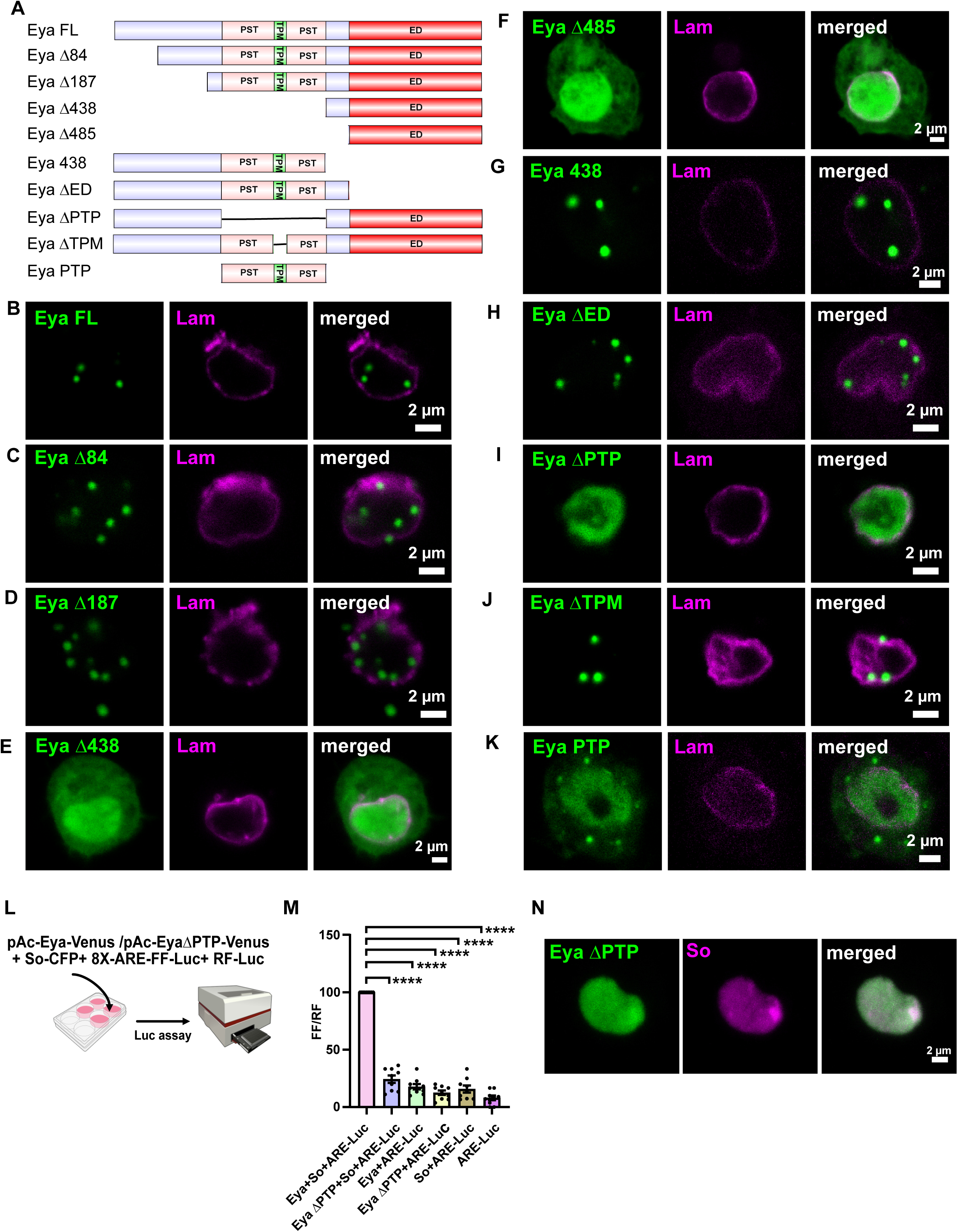
The PST-TPM-PST region of Eya is crucial for condensate formation. **A.** Schematics of all truncated Eya constructs **B-K.** Representative image of S2 cell immunostained with lamin antibody to mark the nucleus and expressing **(B)** full length Eya, **(C)** Δ84 Eya, **(D)** Δ187 Eya, **(E)** Δ438 Eya, **(F)** Δ485 Eya, **(G**) N-terminal 438 amino acid fragment of Eya, **(H)** ΔED Eya (485 Amino acid fragment from N terminal), **(I)** ΔPTP Eya (PST-TPM-PST domain deleted), **(J)** ΔTPM Eya (TPM region deleted), **(K)** PTP fragment of Eya (PST-TPM-PST). **L.** Schematic of the reporter gene-based transcription activation assay. S2 cells were transfected with the 8XARE-Firefly luciferase and Renilla luciferase to normalize the values along with Eya and So, Eya ΔPTP and So, Only Eya, only Eya ΔPTP, and only So. Transfected cells were later lysed and processed for dual luciferase assay using a luminometer. **M.** Quantitation of reporter gene activity normalized with Renilla luciferase activity shows that while the Eya and So can activate the reporter gene transcription, the ΔPTP mutant with So cannot activate the reporter gene transcription. The values for Eya+So were first normalized to 100 and were then used as a reference to normalize the other columns. RM one-way ANOVA with Geisser-Greenhouse correction was performed. The mean values of each column were compared with the mean of the Eya+So column with the help of Dunnett’s multiple comparison test. The p-values of every column compared to Eya +So were <0.0001. The experiment was repeated three times with 3 technical repeats for each time. Error bars represent SEMs. **N.** Representative image of S2 cell coexpressing Eya ΔPTP and So, showing no partitioning of So in condensates in the absence of Eya condensates.

We noticed that three of the deletion constructs Δ438, Δ485, and ΔPTP showed diffused patterns of expression, and the only region common between these three was the PTP region. This suggested the PTP domain is the minimum region that is indispensable for condensate formation by Eya. We also tested this in reverse by imaging a construct encompassing the PTP region alone (Eya PTP) and observed this to form condensates, even though these were not nuclear (Figure 4K), possibly due to the loss of the nuclear localization sequence. We further tested the ability of the PTP region to form condensates in vitro. We expressed this in E.coli with an N-terminal MBP tag and a C-terminal GFP tag with 6X His, with both the MBP and His tags removable by TEV enzyme. We purified this protein and, in conditions mimicking crowding using 10% Dextran, in the presence of TEV, we were able to see the formation of condensates within one hour (Supplementary Figure 4).

Based on all these observations, we infer that the PST-TPM-PST (PTP) region is crucial for condensate formation.

### The PST-TPM-PST region of Eya is needed for transcriptional co-activation with So

We next addressed the consequences of transcription regulation on interfering with the condensate formation. We checked for the ability of Eya and So to act in transcription activation using the previously described 8X-ARE Luciferase construct (Silver *et al*, 2003). In our reporter-based assay, where we monitor the Luciferase activity and normalize with a control Renilla Luciferase activity (Figure 4L), we found that only when both Eya and So are together, we were able to see the activation, but not with the solo expression of Eya and So (Figure 4M). When we replaced the full-length Eya construct with the

ΔPTP domain construct, which does not form condensates, we saw significantly reduced luciferase activity (Figure 4M). This finding is in agreement with the previous report on the role of the PTP region in transcription activation (Silver *et al*, 2003). Eya ΔPTP has been reported to retain the ability to interact with So (Jin & Mardon, 2016), suggesting the observed reduction in transcription activity is not due to the failure of Eya to interact with So but most likely due to the lack of condensate formation of Eya. In cells coexpressing EyaΔPTP and So, we do not see any So as condensate (Figure 4N), suggesting that the partitioning of the transcription factor So by Eya is required for transcription regulation.

Interestingly, the PTP domain identified here as crucial for condensate formation and transcription coactivation was identified as crucial for eye development, as a genomic construct lacking the PTP region is unable to rescue the lack of eye phenotype of the eya^2^ mutant (Jin & Mardon, 2016). Thus, these observations connect the condensate formation capacity of Eya with transcription coactivation and eye development.

### A human disease-associated mutation alters Eya condensate properties and function

Drosophila Eya has four homologs in vertebrates, Eya1-4 (Xu *et al*, 1997; Zimmerman *et al*, 1997; Borsani *et al*, 1999) (Supplementary Figure 5). Vertebrate Eya homologs have been earlier reported to rescue the Drosophila Eya mutant’s lack of eye phenotype (Bui *et al*, 2000; Mutsuddi *et al*, 2005). All the human Eya 1-4 have disordered domains, and GFP-tagged human Eya1-4 (hEya1-4) form condensate-like structures in HEK293T cells (Supplementary Figure 6A, B).

Mutations in human Eya genes are associated with altered development of the ear, kidney, and muscles, and mutations in these are present in several disease conditions, such as Branchiootorenal (BOR) Syndrome, Branchiootic (BO) Syndrome, Deafness, cardiomyopathy(Tadjuidje & Hegde, 2012). We investigated whether the residues associated with such mutations can affect the condensate behavior and transcription regulation. We focused on mutations within the PST-TPM-PST region as we found this region to be crucial for condensate formation.

We first aligned Drosophila Eya and human Eya’s and narrowed down the regions in human Eya’s that are homologous to the PST-TPM-PST domain amino acids of Drosophila Eya (Figure 5A). Next, we identified residues that are mutated in diseased conditions and are also conserved between human and Drosophila. These are hEya1G135S, hEya1G140S and hEya1Y226C (Figure 5A). We introduced these mutations in Drosophila Eya as G223S, G228S, and Y337C and observed that all three mutants form condensates and colocalize with So (Supplementary Figure 7), suggesting their condensate-forming ability and ability to partition So are not abolished in these. We next tested these with So and 8X-ARE-Luciferase in S2 cells for their transcription regulation property. Amongst the three mutations tested, we observed the G223S mutation to show significantly reduced transcriptional activity (Figure 5B). The G223S mutation is homologous to the G135S mutation in the Eya1 gene, which was earlier reported to be associated with deafness (Miyagawa *et al*, 2013). We performed live imaging of the wild-type Eya and G223S expressing cells to observe these condensates. We found that while the wild-type Eya condensates undergo fusion, the G223S mutant condensates, even when in proximity, don’t undergo fusion (Figure 5C). In FRAP experiments, while we noticed rapid recovery in wild-type Eya condensates, the G223S condensates showed no recovery and significantly reduced mobile fraction (Figure 5D, E, F). This suggested that the G223S mutation alters the condensate behavior of Eya, most likely making these condensates more solid-like. Together with the ARE luciferase reporter assay, this suggested that Eya condensates with diminished transcriptional activity are associated with altered condensate behavior.

**Figure 5.**
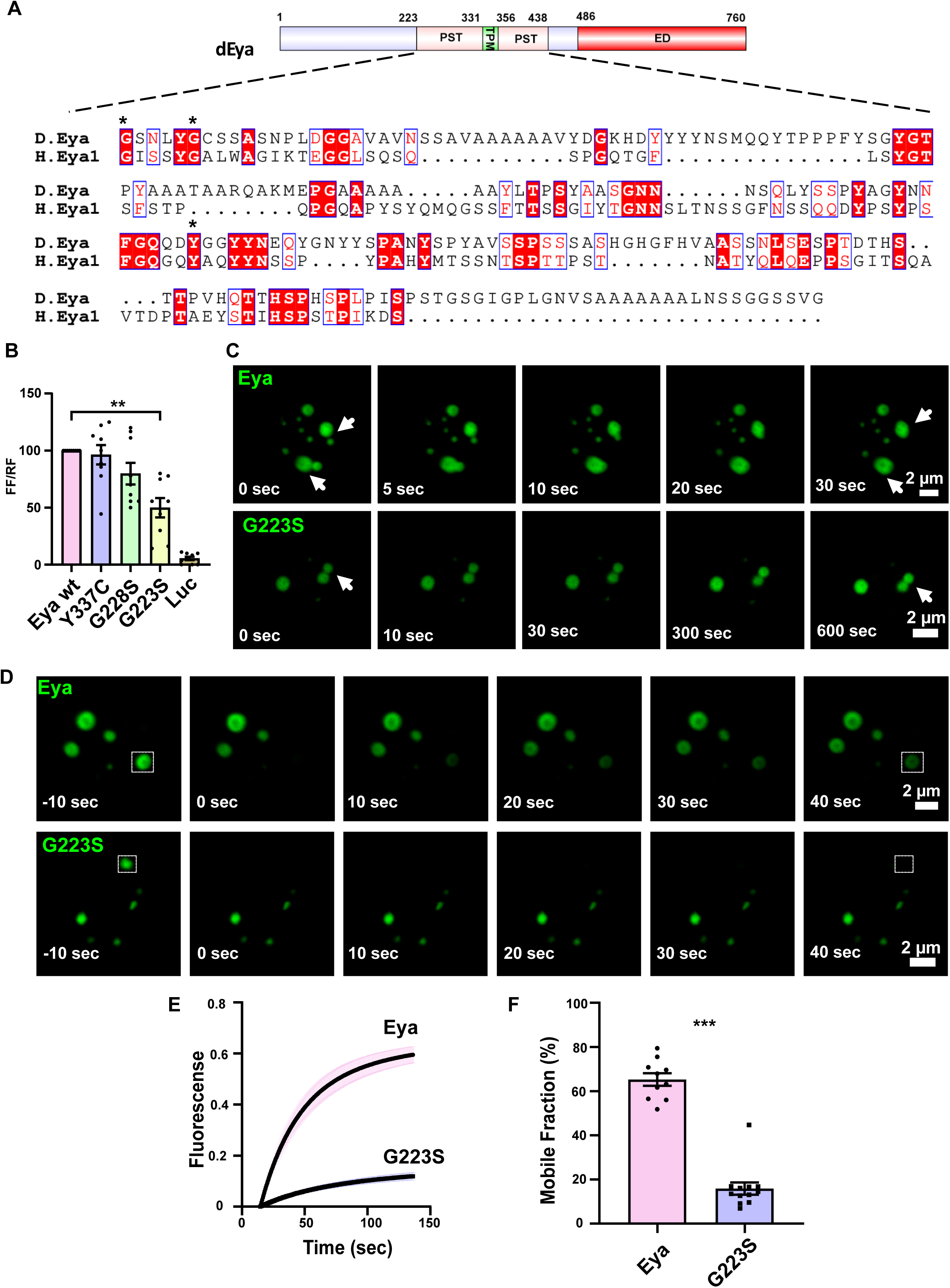
The deafness associated G223S mutation alters the material property of Eya condensates. **A.** Schematic showing alignment of the PST-TPM-PST region of Drosophila Eya with the corresponding region from Human Eya1. The residues inside the red boxes are residues associated with mutations in human patients. **B.** Quantitation of reporter gene activity normalized with Renilla luciferase activity for wild type and the three Eya mutants shows the G223S mutation to exhibit significantly reduced reporter gene activity. The values for Eya wt were normalized to 100. RM one-way ANOVA with Geisser-Greenhouse correction was performed, while the mean difference was calculated using Dunnett’s multiple comparison test, where Eya wt was considered as the reference point. The p-value for the G223S mutant was 0.0085. Error bars represent SEMs. **C.** Frames from live confocal imaging of S2 cells expressing wild-type Eya (top panel) showing fusion of the condensates (marked by arrow), whereas the Eya G223S mutant (bottom panel) condensates do not fuse. **D.** Frames from the FRAP experiment with wild-type Eya condensates (top panel) and G223S (bottom panel) showing the state of the recovery of the fluorescence after photobleaching (marked by a white box at the beginning and the end). **E.** FRAP recovery graph of wild-type Eya condensate and the G223S mutant. **F.** Quantitation shows G223S mutant condensates to have significantly reduced mobile fraction in comparison to the wild-type Eya. Unpaired Student’s t-test was done to compare the difference between mobile fractions. N for Eya wt =10 and for G223S =12. The p-value was <0.0001. Error bars represent SEMs.

**Figure 6.**
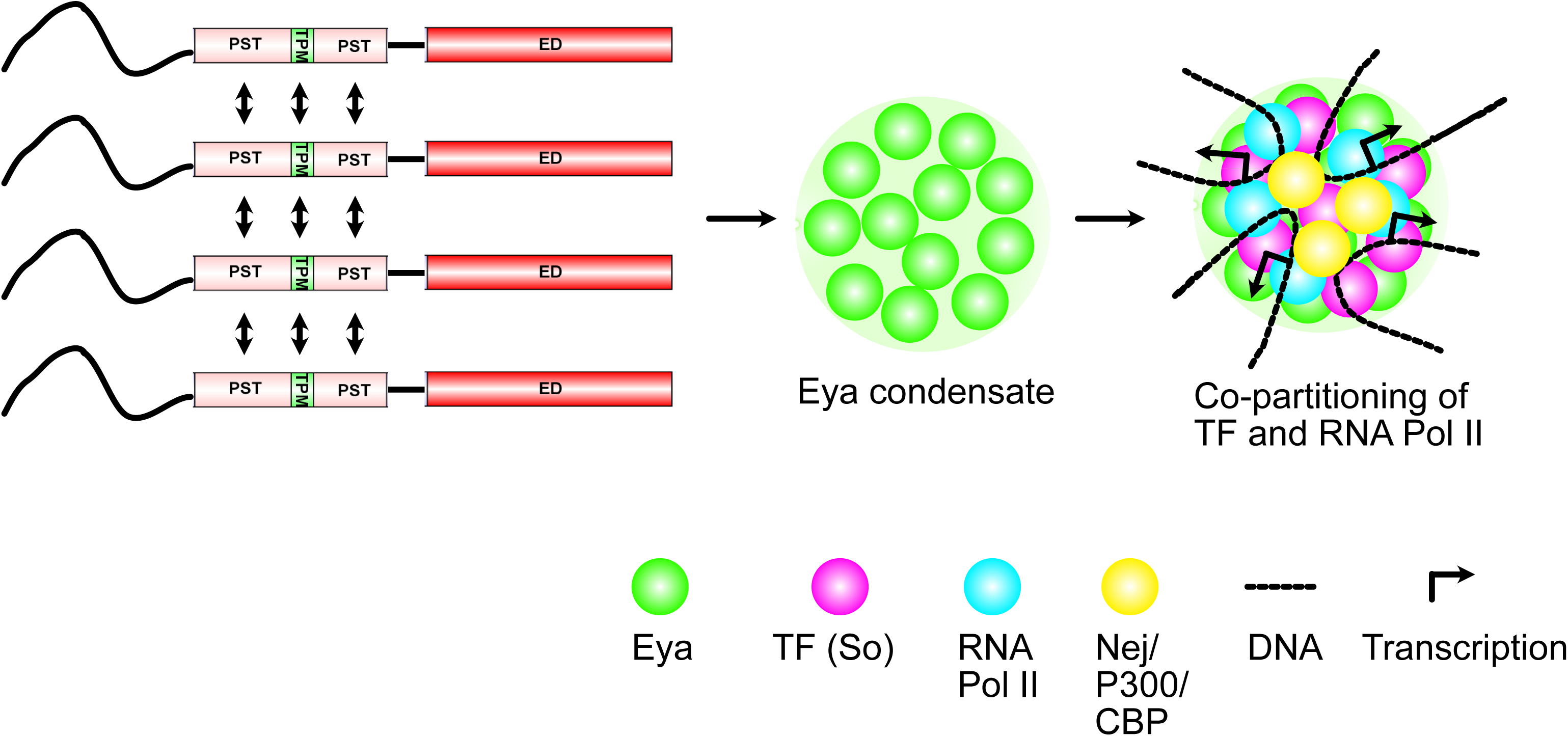
Schematic of the suggested model of Eya-centric transcription activation. The PST-TPM-PST (PTP) region of Eya interacts amongst themselves to form the Eya condensates, which can then partition the transcription factor (So) and RNA pol II in them and activate transcription.

## Discussion

Eya forms nuclear condensates that fuse amongst themselves and, on photobleaching, show very rapid recovery. The equilibrium of Eya between the condensate and non-condensate diffused state can be shifted by using osmotic challenges, where hypotonic treatment shifts towards the diffused state and hypertonic treatment shifts them to form a condensate state. This osmolarity-mediated modulation of condensate property is a shared property for several other condensate-forming proteins. The viral protein MxA (Davis *et al*, 2019; Sehgal *et al*, 2022), Dcp1a in processing bodies and stress granules (Jalihal *et al*, 2020), kinases Ask3 and Wink (Watanabe *et al*, 2021; Boyd-Shiwarski *et al*, 2022; Morishita *et al*, 2023), ALS associated Fus and Tdp-43 proteins (Gao *et al*, 2022), regulators of Hippo pathway AMOT and KIBRA (Wang *et al*, 2022a) and the transcriptional coactivator YAP (Cai *et al*, 2019) have all been reported to undergo osmotic treatment based remodeling. Together, the fusion behavior, rapid recovery on photobleaching, dissolution of the condensates by 1,6-HD, and osmotic remodeling suggest the Eya condensates to be liquid-like in nature.

Eya condensates cause selective co-partitioning of So, Optix, Ey, Dac, and Nemo proteins. This partitioning is not dependent on the presence of intrinsically disordered regions, as even the proteins not showing co-partitioning, Abd-A, Abd-B, Tim, and Socs44A, have intrinsically disordered regions, whereas Nemo, which partitions, does not have disordered regions, suggesting the co-partitioning has some specificity connected with associated function (Supplementary Figure 8). So, Optix and Ey are homeobox-containing transcription factors, and Dac contains a helix-turn-helix family, and all these proteins can bind to DNA. Because Eya alone cannot bind to DNA directly, through the selective partitioning of its interactors with DNA binding ability, which are the transcription factor members of the retina determination gene network pathway, it can create selective spatial zones or transcription hubs where the promoters of target loci can be enriched and transcribed. This model gets further support due to the presence of Nej/P300/CBP, the RNA PolII machinery, acetylated H2B, a marker for active enhancers, and RNA within the Eya condensates.

Our findings align with broader evidence that transcriptional regulation involves phase separation or condensate formation of co-activators and transcription factors. Liquid-like condensates of coactivators BRD4 and MED1 were found to concentrate RNA Pol II on super-enhancers (Sabari *et al*, 2018). The transcription factors, Oct4, Myc, P53, Nanog, Sox2, RARa, GATA2, Estrogen receptor, and GCN4 also showed partitioning in Med1 and Med15 coactivator condensates (Boija *et al*, 2018). The coactivator P300/CBP forms condensates and uses its disordered domain to co-condense with the transcription factors Sox2 and p65/NFKB, promoting the transactivation of P300 and further Histone acetylation, recruitment of Brd4, and increasing the transcriptional activation (Ma *et al*, 2021). PGC1-A, a transcriptional coactivator, has been found to form nuclear condensates of a liquid-like nature through its disordered C-terminal domain, which, through its RNA binding, assembles other protein complexes involved in transcription regulation (Pérez-Schindler *et al*, 2021). Myocardin, a transcription coactivator lacking DNA binding ability, forms condensates through its C-terminal disordered domain, partitions DNA binding transcription factors SRF and myocyte enhancer factor 2C along with RNA Pol II, to activate the transcription program for smooth muscle cells and cardiomyocyte differentiation from human induced pluripotent cells (Gan *et al*, 2024).

Similar to these, we find the Eya co-activator condensates to partition the transcription factors So, Dac, Optix, and Ey. With our focus on Eya and So, we observe these condensates to partition RNA pol II and suggest these sites to be the sites for transcription of its reporter (Figure 7). In the context of the RDGN pathway, Eya is connected with Ey, So, Dac, and Optix in a multilayered regulatory manner with positive and negative feedback (Kumar, 2010; Davis & Rebay, 2017). At the beginning of the development of eye discs, Ey is expressed throughout the disc and induces the expression of Eya and So, which starts the differentiation process. Eya and So feedbacks to induce the expression of Ey (Bonini *et al*, 1997; Halder *et al*, 1998; Niimi *et al*, 1999; Pignoni *et al*, 1997; Pauli *et al*, 2005). On the contrary, in the differentiated cells and the morphogenetic furrow (MF), the expression of Ey needs to be suppressed, and this is done by Eya and So (Atkins *et al*, 2013). This suggests depending on the context, the same Eya and So complex can be an activator and also switch to a transcription repressor. Eya and So together regulate the expression of Dac and Optix, and both these can feedback to Eya and So (Li *et al*, 2013b; Pignoni *et al*, 1997). While our work suggests the Eya condensates partitioning So can be sites for transcription activation, one of the drawbacks of the work is that, in the absence of information on direct DNA binding targets of the other partitioned transcription factor proteins, we have not been able to test what role the Eya condensates can play when Dac, Optix, or Ey is partitioned in them. The mechanism of how the Eya, So complex, turns from activator to repressor is also not clear. Based on findings of mammalian homolog of Drosophila Dac, Dach1 acting as a repressor (Zhou *et al*, 2010a, 2010b; Chen *et al*, 2013; Chu *et al*, 2014), one possibility is that partitioning of Dac in the Eya condensates can turn it into a repressor. It is also possible that the composition and quantity of DNA-binding transcription factors in the Eya condensates change stoichiometrically over time and space of the eye disc, and such a mechanism might help in dosage-dependent transcription regulation during eye development.

The other protein we noted to be sequestered in the Eya condensates is the kinase Nemo. While Nemo mutants show no change in Eya levels, it can genetically interact with Eya to enhance the ectopic eye formation phenotype (Braid & Verheyen, 2008). It is possible that the recruitment of Nemo in the Eya condensates can phosphorylate some of the components in the condensate, including Eya, which was earlier identified as a direct substrate of Nemo, and this phosphorylation can positively regulate the transcription process and amplify the output of Eya and So mediated transcription activation. Also, one of the known substrates of Nemo is Mad (Mothers against Dpp), a component of the TGF-β/ Dpp/ BMP signaling pathway (Zeng *et al*, 2007; Merino *et al*, 2009). Dpp signaling is closely connected with eye development, as evidenced from Dpp mutants showing reduced expression of Eya and So, and Dpp signaling is reduced in Eya mutants (Hazelett *et al*, 1998; Chen *et al*, 1999; Weasner & Kumar, 2022). One possibility is that the partitioning of Nemo in Eya condensates can regulate the integration of the Dpp signaling pathway with the retinal development pathway. In mammalian cells, the occurrence of such a possibility integrating signaling pathways with condensates was demonstrated when terminal signaling factors of the WNT, JAK/STAT, and TGF-β pathways were shown to partition in mediator condensates on super-enhancers to regulate transcription (Zamudio *et al*, 2019).

Our deletion mapping experiments suggested the PTP region is crucial for Eya’s condensate formation and transcription regulation. The ΔPTP mutant doesn’t form condensate. Eya has an ED domain in its C-terminal through which it interacts with So (Pignoni *et al*, 1997). In the absence of the PTP region, the ΔPTP mutant can interact with So (Jin & Mardon, 2016), possibly through the intact ED domain. However, in our reporter-based assay to measure transcription, we notice low levels of Luciferase expression with the ΔPTP mutant in comparison to the full-length Eya control, suggesting that in the absence of Eya condensates, the transcription factor So cannot be partitioned and hence causes significantly reduced activity. As the same region was previously shown to be the domain indispensable for eye formation, our findings link a possible connection between condensate formation, transcription regulation, and eye development. The PTP region is referred to as the transactivation domain, and this is in agreement with observations of other transactivation domain’s in Myc (Yang *et al*, 2024), Yin Yang1 (YY1) (Wang *et al*, 2022b), and Lymphoid enhancer factor 1 (LEF1) (Zhao *et al*, 2023), importance in condensate formation in transcription regulation.

We further screen for mutations in the PTP region of Drosophila Eya at conserved sites with respect to its human homologs, which have been reported to be associated with disease conditions. Here, we identify the G223S mutation associated with deafness to change the material property of the Eya condensates by shifting them from a liquid-like to a solid-like state, as evidenced by the live imaging and FRAP experiments. A consequence of the shift of the material property is that the condensates can still recruit So, but this mutant Eya shows significantly lesser co-activation of the reporter gene. Also, the G to S mutation changing the material property is supported by earlier work trying to decipher the grammar behind phase separation, suggesting that Glycine helps in maintaining the condensates in a liquid-like state, whereas Serine pushes them towards a more hardened state (Wang *et al*, 2018). The Glycine to Serine transition is also reported to cause dynamical arrest in condensates (Alshareedah *et al*, 2024). Alternatively, the G to S mutation can cause a change in the interface properties of the condensates, which interferes with the fusion and dynamicity of the condensates.

Previously, Drosophila Eya has been shown to be needed for the active maintenance of neuronal identity post-differentiation (Eade *et al*, 2012), but the mechanism is not clear. Post differentiation, the maintenance of the differentiated identity is suggested to be an active process through continuous maintenance of the actively transcribing state (Blau & Baltimore, 1991; Leyva-Díaz & Hobert, 2019; Deneris & Hobert, 2014). Phase-separated condensates through trapping the transcription machinery in transcription sites are suggested to play a role in this (Gurdon *et al*, 2020; Javed & Gurdon, 2022; Javed *et al*, 2022). We suggest, transcription regulation by Eya through condensate formation can be a possible mechanism for this.

In summary, our study connects the liquid-like condensate property of the coactivator Eya with the regulation of transcription co-activation in developmental processes and provides mechanistic insight into how disease-associated mutations in Eya might disrupt gene regulation by altering condensate dynamics.

## Materials and Methods

### Constructs

The Eya constructs were made by cloning the Eya ORF without the stop codon in Topo-D-Entr followed by LR cloning in pAWV vectors. Truncated constructs were also made similarly. The primer sequences are given below:

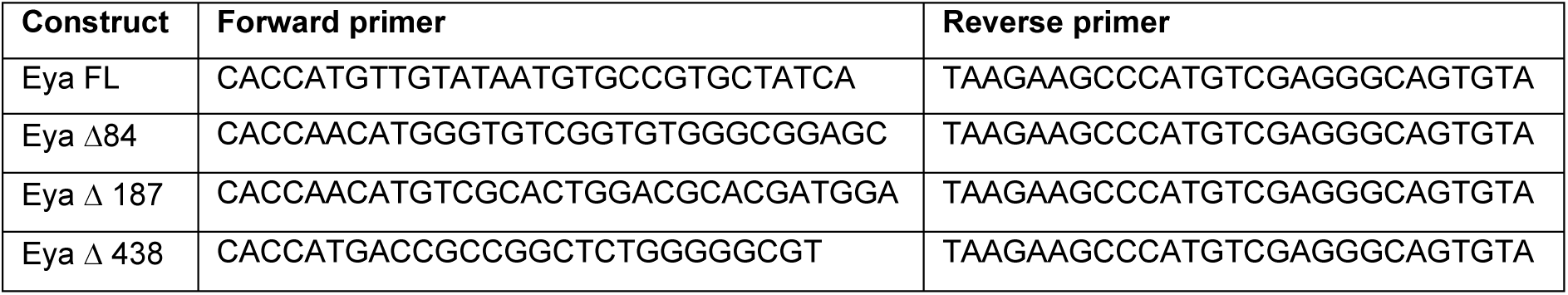
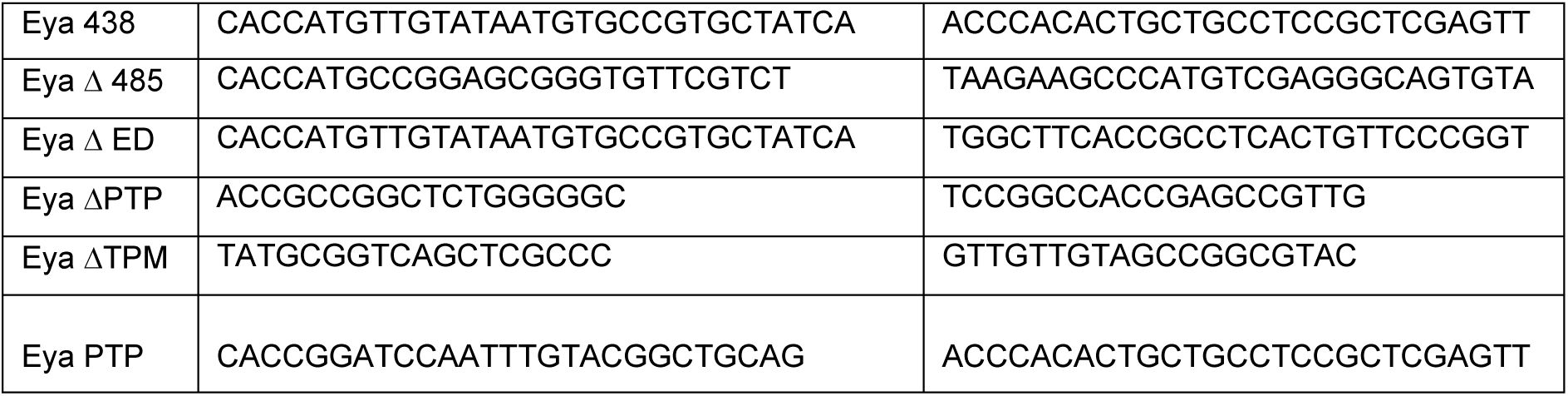

The Gateway Entry clones for Eya interactor constructs were purchased from the FlyBiORFeome collection of DGRC and LR cloned into the pAWC destination vector. The following is the list of Gateway Entry clones from DGRC that are used in this study:

| Name of gene | Clone ID |
| --- | --- |
| Abdominal A (Abd-A) | GEO12744 |
| Abdominal B (Abd-B) | GEO02093 |
| IKK-Beta | GEO02223 |
| Hop | GEO12249 |
| Relish (Rel) | GEO04678 |
| Timeless (Tim) | GEO01888 |
| Socs44A | GEO09041 |
| Optix | GEO03888 |
| Eyless (Ey) | GEO09847 |
| Lamin C (LamC) | URS01250 |
| Fibrillarin | GEO13932 |

Constructs for expressing GFP-tagged human Eya proteins (hEya1-4), CFP-tagged Dac and Nemo, and mRFP-tagged Rpb3 were gene-synthesized from Twist Bioscience. The pCasper4-His2Av-RFP and 8X-ARE-Luciferase constructs were kind gifts from Stephan Heidmann and Francesca Pignoni and are previously described (Silver *et al*, 2003; Schuh *et al*, 2007; Kenyon *et al*, 2005b).

The constructs were transfected in Drosophila S2 cells using Effectene (Qiagen) as per the manufacturer-recommended protocol. hEya1-4 GFP-tagged constructs were transfected into HEK293T cells using Lipofectamine.

### Fly stocks

The flies were raised on a standard cornmeal food and kept in a 22℃ incubator with a 12:12 hr day, night cycle. The following is the list of fly lines that are used in this study.

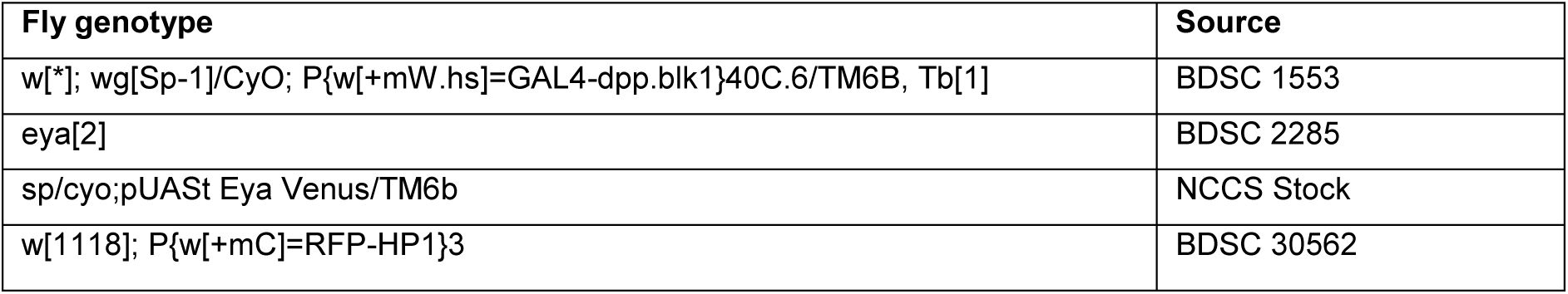
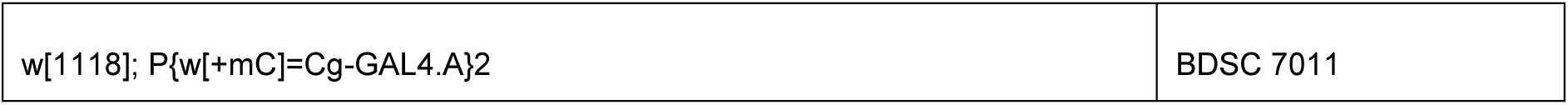

### Imaging and FRAP analysis

Imaging and FRAP experiments on the transfected cells were performed on a Nikon A1R confocal microscope. During imaging, saturation of the condensates was avoided by controlling the laser power and gain according to the expression levels in the cells. FRAP movies were analyzed using a Jython script as described here (https://imagej.net/tutorials/analyze-frap-movies). For imaging of two colors, a sequential mode of imaging was used to avoid any bleed-through problems. For the smiFISH and Rpb3 co-imaging experiment, the cells also contained So-CFP, but were not imaged along with the other two colors.

### Immunostaining

S2 cells expressing Eya-Venus were taken out on a coverslip-bottom dish that was coated with Concanavalin A. The cells were fixed with 4% PFA for 10 minutes, washed thrice with 1x PBST (1x PBS + 0.1% Triton-X-100), and blocked with 5% Normal Goat Serum (NGS) diluted in 1x PBST for 1 hr. The primary antibody diluted in blocking solution was added next and incubated at room temperature for 2 hrs. Post removal of primary antibody, the cells were washed thrice with 1x PBST and incubated with fluorophore-tagged secondary antibody diluted in blocking solution for 1 hour. After the removal of the secondary antibody and 3x washes with 1x PBST, the cells were imaged in a confocal microscope. For immunostaining Drosophila tissues, eye imaginal discs from wandering third instar larvae and ovaries from 5-day-old adult egg-laying females were dissected out in dissecting buffer (0.1M phosphate Buffer) and then fixed with 2% Paraformaldehyde-Lysine-Periodate (PLP) solution for 30 minutes. After removing the fixative, the tissues were washed thrice with Wash Buffer (0.1M Phosphate Buffer 0.1% Triton-X-100) and then put in blocking solution (5% NGS in 1xPBST) for 2 hrs at RT. Next, the tissues were incubated overnight with the primary antibody at 4℃. On the following day, the primary antibody was removed, and tissues were washed thrice with Wash Buffer before the addition of the secondary antibody. After 2hr incubation with secondary antibody at RT, the tissues were again treated with 3x Wash Buffer. The tissues were next counterstained with Phalloidin to mark the cell boundary and Hoechst to mark the nucleus. Both the dyes were dissolved in 1xPBS and incubated for 15 minutes each. A final 1xPBS wash was given before mounting the tissues on a slide with a drop of Vectashield in between the slide and coverslip.

### Reporter gene assay for transcription

The 8X-ARE luciferase construct (kind gift from Dr. Francesca Pignoni), along with a pAc-Renilla Luciferase construct, was transfected with Eya Venus and So-CFP. Cells were grown for 48 hours, followed by pelleting them down for the Luciferase assay using the Biotium Firefly & Renilla Luciferase Single Tube Assay Kit and taking the readings in a luminometer.

### smiFISH

smiFISH experiments were performed as per protocols described in (Tsanov *et al*, 2016). Imaging of these experiments was performed using Airy scan mode in a Zeiss LSM 880 confocal microscope.

### 1,6-Hexanediol experiment

1,6-HD was directly dissolved in S2 cell media to make a 5% (v/v) solution. S2 cells expressing Eya FL-Venus 48 hours post-transfection were plated on a concanavalin-A-coated imaging dish. After 15 minutes, the media was carefully removed without disturbing the cells, and 5% (v/v) 1,6-HD solution was added to the dish. After 5 minutes of treatment, 1,6 HD was removed, and cells were washed briefly with 1x PBS. Next, the cells were fixed with 4% para-formaldehyde for 10 minutes, followed by washes with 1xPBS thrice for 10 minutes each. Z-stacked images were taken using a Nikon A1R confocal microscope. The number of puncta per cell was counted manually from the maximum intensity projection of individual images.

### Osmotic challenge experiments

For hypertonic treatment, sorbitol was dissolved in S2 media at a final concentration of 1 𝜇g/ml. For the hypotonic experiment, cells were treated with hypotonic buffer (HEPES-KOH 10mM, pH 7.6, MgCl_2_ 1.5mM, KCl 10mM, and DTT 0.5mM). Both treatments were performed on 48-hour post-transfected S2 cells expressing Eya FL-Venus and So-CFP.

### Antibodies and other reagents

The following is the list of commercially available antibodies and chemicals used in the paper:

| Name of antibody (Dilution used) | Catalogue No. |
| --- | --- |
| Anti-Lamin (1:500) | ADL67.10-s (DSHB) |
| Anti-GFP (1:800) | MBL 598 |
| Anti-acetyl Histone H2B antibody | Sigma 07373 |
| Alexa Fluor 555 anti- mouse (1:1000) | A21424 (Life Technologies) |
| Alexa Fluor 555 phalloidin (1:1000) | A34055 (Life Technologies) |
| Alexa Fluor 488 anti-Rabbit (1:1000) | A21429 (Life Technologies) |
| Hoechst | Invitrogen |
| 1,6-Hexanediol 240117 | Sigma Aldrich |

### Site-Directed Mutagenesis

Disease-related point mutations in the human Eya 1 gene (hEya1) were selected from the public database, ClinVar. Mutations that fell in the PST-TPM-PST region of Human Eya were selected and then traced back to the Drosophila Eya sequence through a ClustalW alignment. The following are the disease-associated point mutations of hEya1 and their Drosophila homologs that are generated in this paper:

| Mutation in hEya1 | Reference | Corresponding mutation in Drosophila Eya |
| --- | --- | --- |
| Y226C | cbioportal database | Y337C |
| G135S | (Miyagawa *et al*, 2013) | G223S |
| G140S | HGMD database | G228S |

SDMs to create single amino-acid substitutions in Eya-Venus were done with the help of a Q5 Site-Directed Mutagenesis kit from NEB. The following primers were used to create the single-point mutation in the Eya-pAWV plasmid.

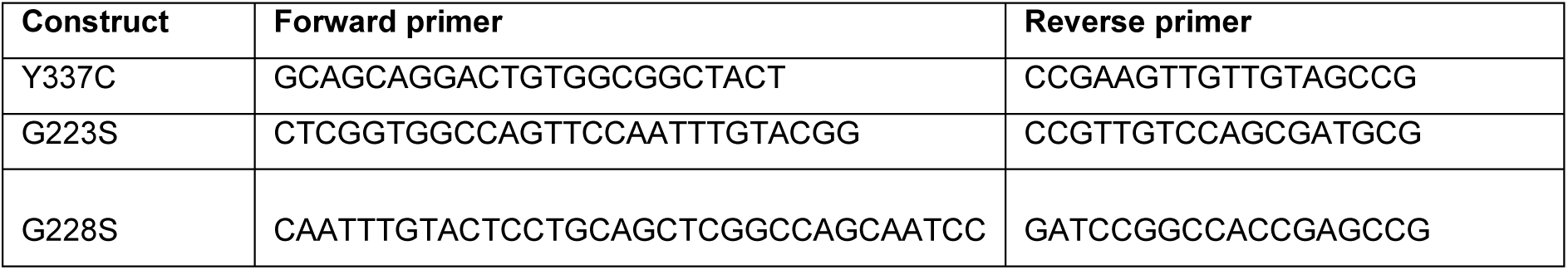

### Eya PTP cloning for protein expression and purification

For protein expression, Eya PTP was cloned into a pMAL-based vector bearing an MBP-TEV tag at the N-terminal and EGFP-TEV-6xHis tag at the C-terminal. At first, primers were designed against Eya PTP such that Sal-I and BamH-I restriction cut sites are incorporated at the 5’ and 3’ ends, respectively. Next, the plasmid vector was double-digested twice; first with SalI and HindIII to get an empty vector backbone and then with BamHI and HindIII to get the EGFP-His fragment. Eya PTP PCR product was digested with SalI and BamHI to generate sticky ends. All three sticky-ended fragments were gel-purified and put in a tri-molecular ligation reaction with DNA ligase overnight at 16°C. The next day, the resultant mixture was transformed into E.coli DH5-𝛼 competent cells. Positive recombinant clones were screened by restriction enzyme digestion and PCR. The selected positive clone was later transformed into E. coli BL21-RIL cells for large-scale protein production.

### Recombinant Protein Purification

For the Drosophila Eya-PTP protein purification, the bacterial culture of BL21-RIL cells containing the MBP and GFP His-tagged construct was induced with 1mM IPTG for 16 hr at 16℃ before harvesting. Next, the cells were lysed using lysis buffer (50 mM NaH_2_PO_4_/Na_2_HPO_4_, 300 mM NaCl, pH=8). The lysate was then incubated with Ni-NTA beads along with 5 mM Imidazole for 2 hrs, followed by washing with wash buffer containing 50mM Imidazole. The final protein was eluted using an elution buffer containing 300 mM imidazole. The fractions having the highest protein concentration were pulled together and dialyzed against storage buffer (50 mM NaH_2_PO_4_/Na_2_HPO_4_, 300 mM NaCl, 10% glycerol, pH 7.5).

### In vitro Phase separation assay

The reaction was set up with TEV enzyme to cleave the MBP tag and with 10% dextran as a crowding reagent. As a control, another reaction was set up without the TEV enzyme. After 1 hour of reaction, images were acquired using a 100× oil lens with 3× zoom.

### Statistical methods

For measuring the significant difference between mobile fractions after FRAP assays, the unpaired Student’s t-test was performed. For analyzing the difference between transcriptional activity in luciferase assays, Ordinary One-way ANOVA was performed. To understand the difference in the number of condensates per cell, unpaired Student’s t-test was performed.

## Acknowledgments

The AM lab was supported by SERB grant CRG/2021/003130, CEFIPRA grant 6503-1, and NCCS intramural funding to AM. We acknowledge Dr. Francesca Pignoni for the 8X-ARE-Luciferase reporter construct, Dr. Stephan Heidmann for the Histone RFP construct, Drosophila Genomics Resource Center, supported by NIH grant 2P40OD010949 for the AWV and AWC and several interactor plasmids, Developmental Studies Hybridoma Bank maintained at the University of iowa for the anti-lamin antibody, Bloomington Drosophila Stock Center (NIH P40OD018537) for several fly stocks used in this study, NCCS Imaging facility for access to the microscopy systems, Maitheli Sarkar and Anamitra Sen for participating in the experimental work at different stages of this work, Dr. Nikhil Ghate for the anti-acetyl H2B antibody, Dr. Rohit Joshi for anti-Nej antibody, Dr. Deepa Subramanyam and Dr. Jomon Joseph for the mammalian cell transfections, Mainak Bose, Girish Ratnaparkhi, Girish Deshpande, Ambuja Navalkar and Justin Kumar for comments on the manuscript. AM thanks Dr. Florence Besse for hosting him in her lab for learning smiFISH and for discussions and comments on the work.

## Author contributions

Conceptualization: AM, ND, JB

Methodology: ND, JB, AM

Investigation: ND, JB

Supervision: AM

Writing—original draft: AM

Writing—review & editing: ND, JB, AM

## Supplemental Data

**Supplementary Video 1.** Live imaging video of S2 cell expressing Eya-Venus showing condensate fusion. The arrow shows the condensates undergoing fusion. The video is taken with imaging frames 10 seconds apart and shown at a 1fps rate.

**Supplementary Video 2.** FRAP imaging video of S2 cell expressing Eya-Venus showing condensate recovery after photobleaching. The box shows the photobleached area and * marks condensate fusion in the cell. The video is taken with imaging frames 10 seconds apart and shown at a 1fps rate.

## Supplemental Figure legends

**Supplementary Figure 1.**
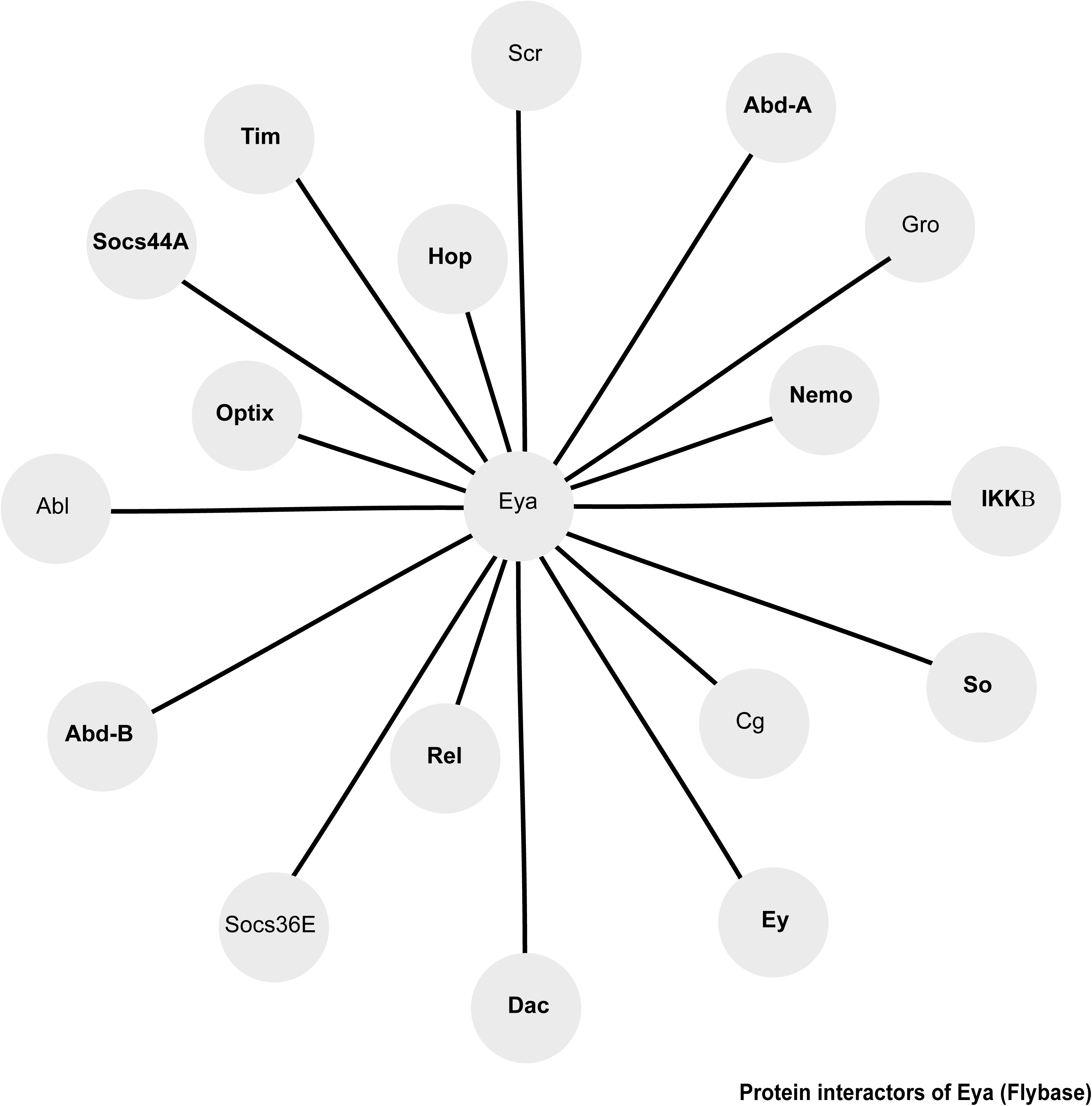
Protein interactor map of Drosophila Eya (adapted from Flybase). Candidate genes used in this manuscript are in bold lettering.

**Supplementary Figure 2.**
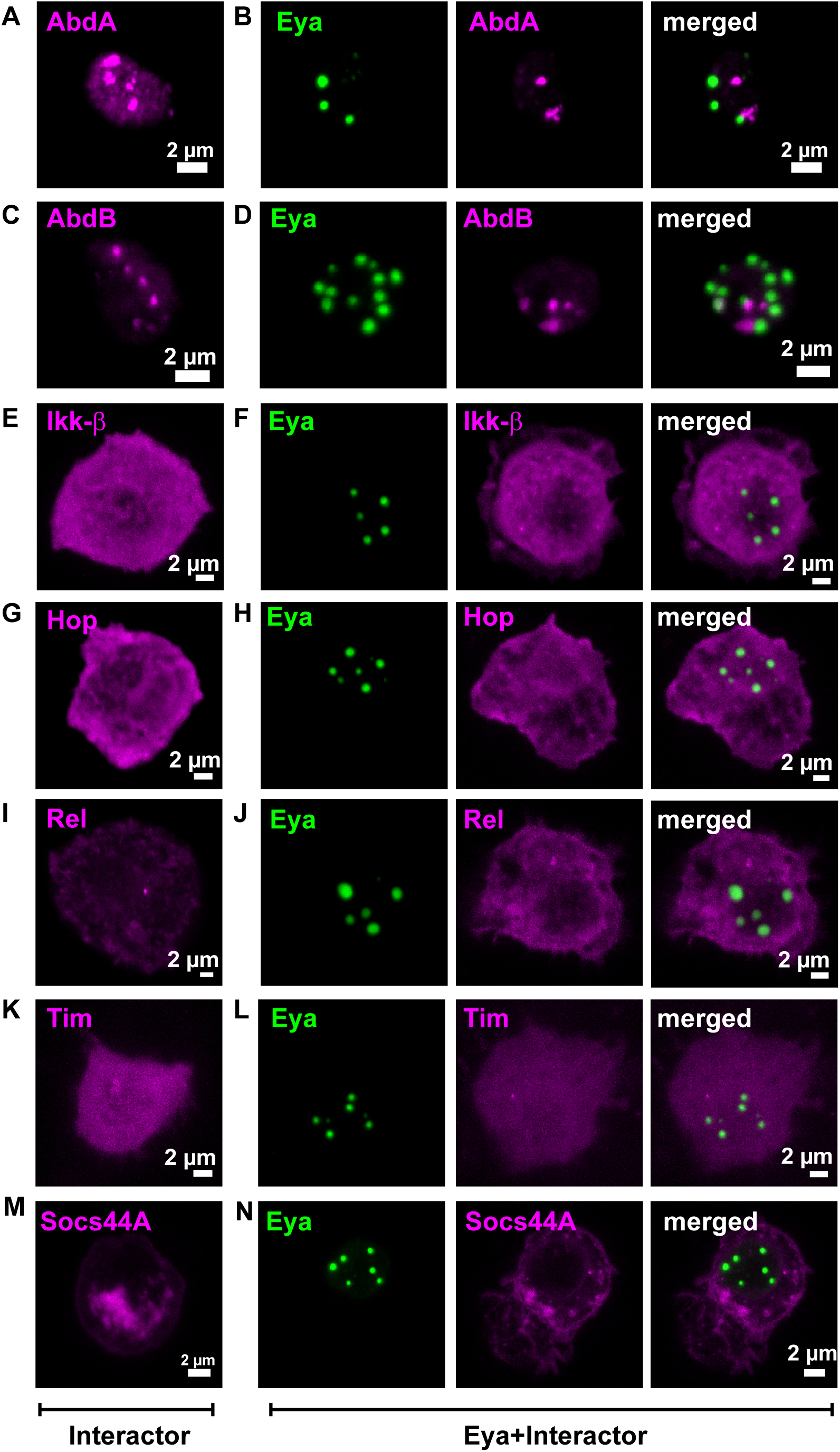
Eya condensates with interacting proteins. **A-N.** Representative image of S2 cell expressing **(A)** Abd-A, **(B)** Abd-A and Eya, **(C)** Abd-B, **(D)** Abd-B and Eya, **(E)** IKK-B, **(F)** IKK-B and Eya, **(G)** Hop, **(H)** Hop and Eya, **(I)** Rel, **(J)** Rel and Eya, **(K)** Tim, **(L)** Tim and Eya, **(M)** Socs44A, **(N)** Socs44A and Eya

**Supplementary Figure 3.**
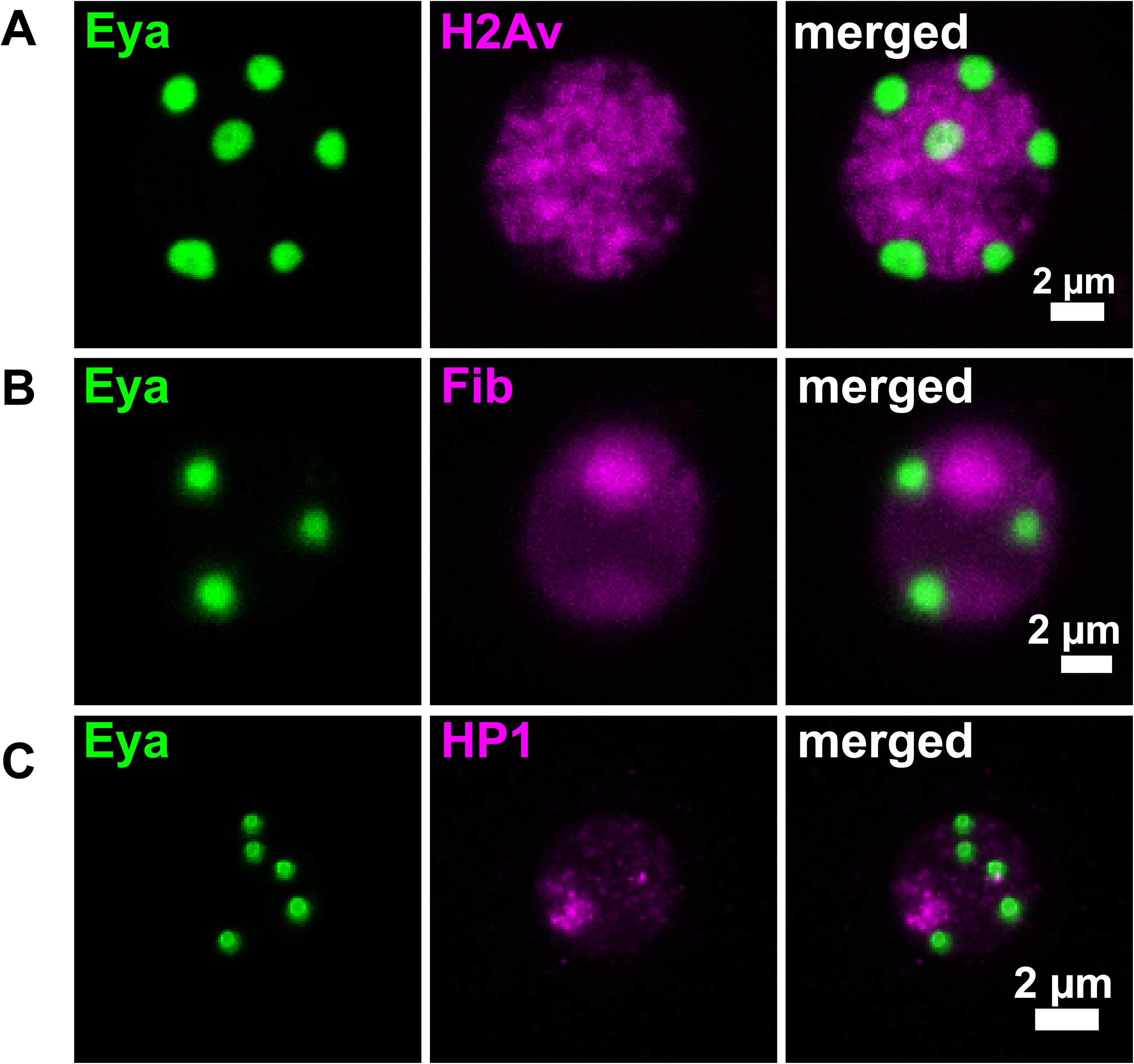
**A.** Representative image of S2 cell coexpressing Eya and Histone2Av showing Eya condensate regions do not colocalize with Histone. **B.** Representative image of S2 cell coexpressing Eya and Fibrillarin showing Eya condensates do not colocalize with Fibrillarin. **C.** Representative image of a cell coexpressing Eya and heterochromatin marker HP1 showing Eya condensates do not colocalize with HP1.

**Supplementary Figure 4.**
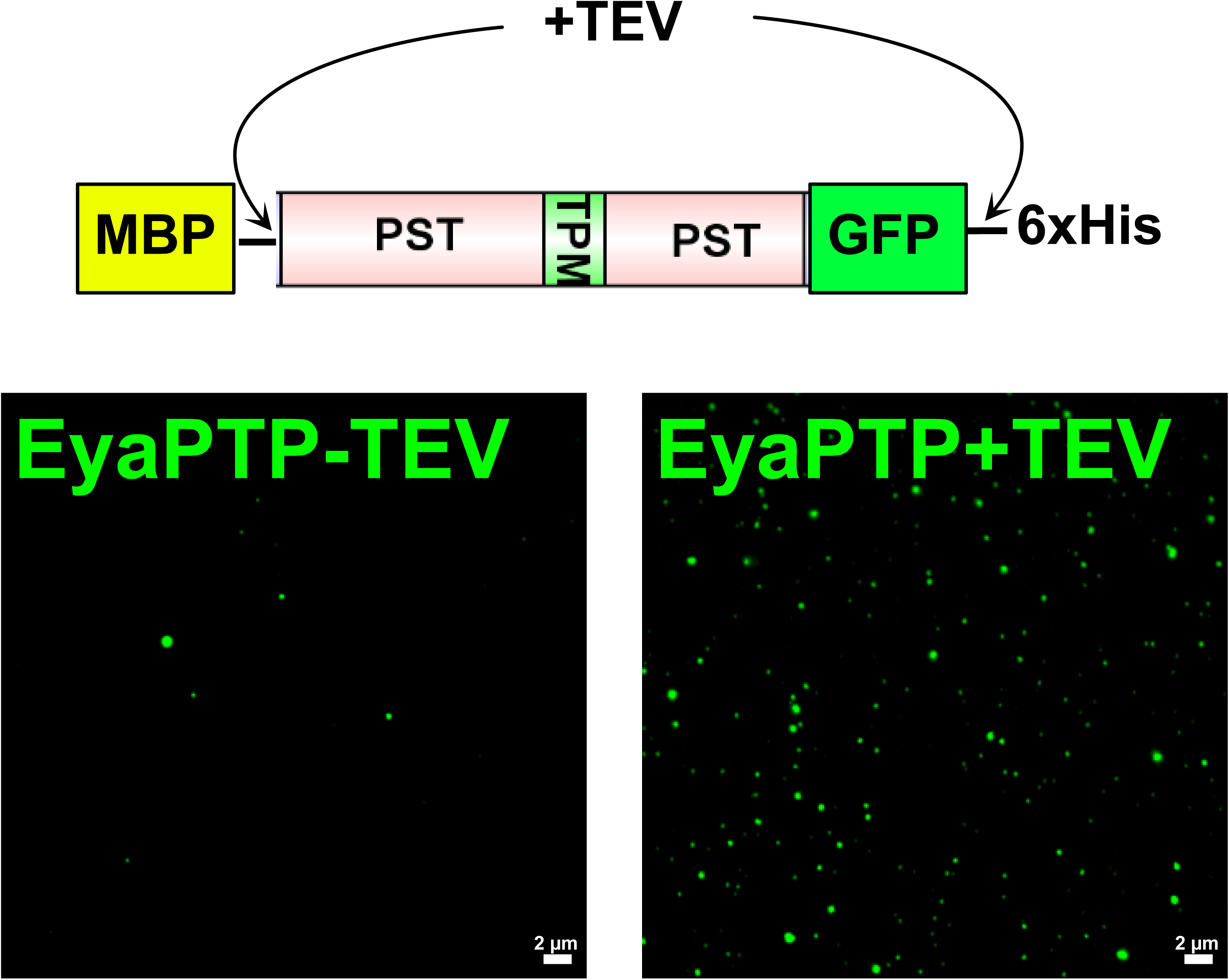
Purified Drosophila PST-TPM-PST domain with N-terminal MBP tag and C-terminal GFP-His tag shows condensate-like structures in vitro on cleaving off the MBP and His tag using TEV.

**Supplementary Figure 5.**
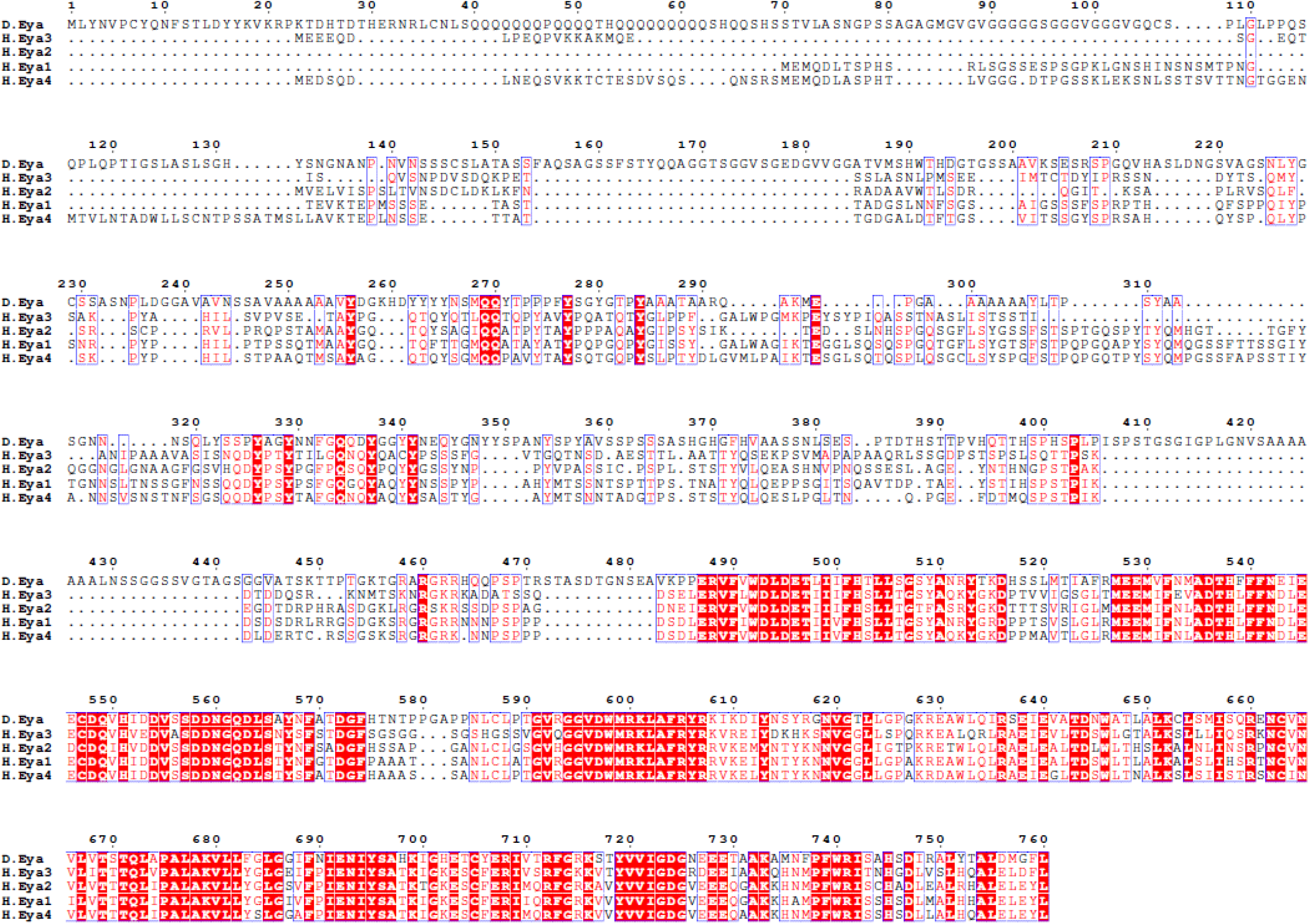
ClustalW alignment of Drosophila Eya with Human Eya1-4.

**Supplementary Figure 6.**
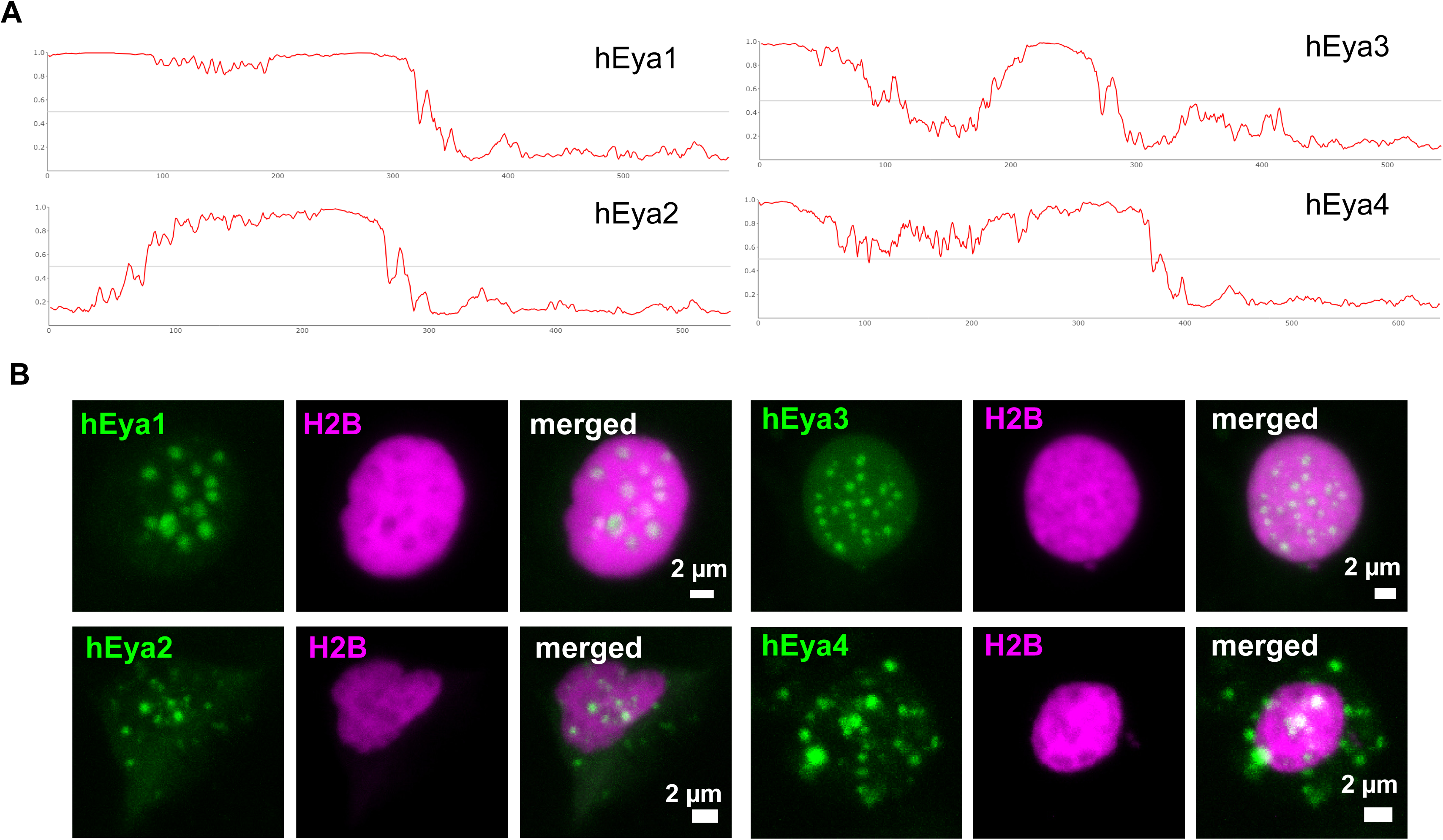
**A.** Intrinsically disordered regions in Human Eya1-4 as predicted using IUPRED3. **B.** GFP-tagged Human Eya1, 2, 3, 4 form condensate-like structures in HEK293T cells

**Supplementary Figure 7.**
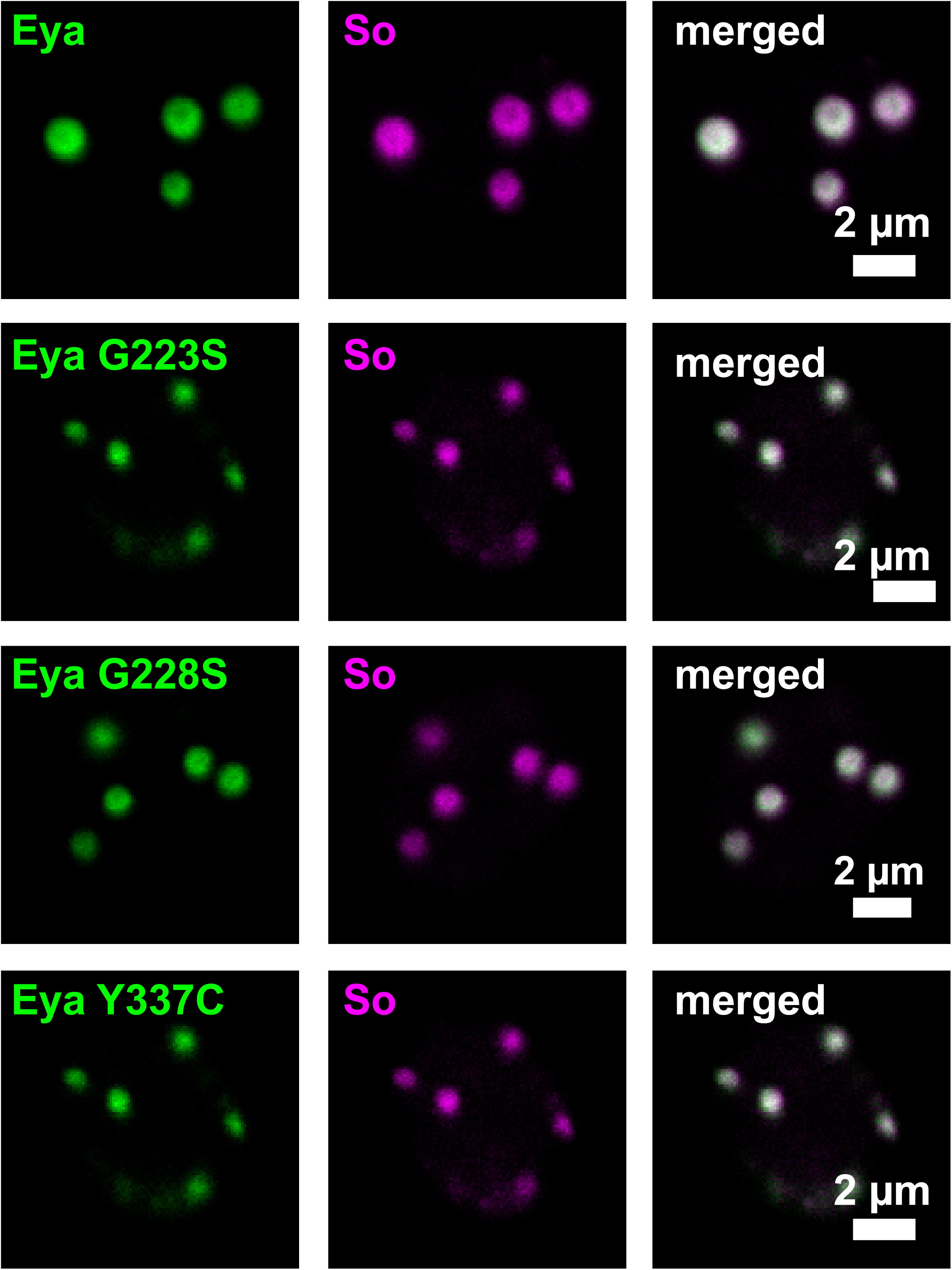
Wildtype Eya and the mutants (G223S, G228S, and Y337C) all form condensates that can copartition So in them, suggesting these mutations do not disturb the co-partitioning ability of Eya.

**Supplementary Figure 8.**
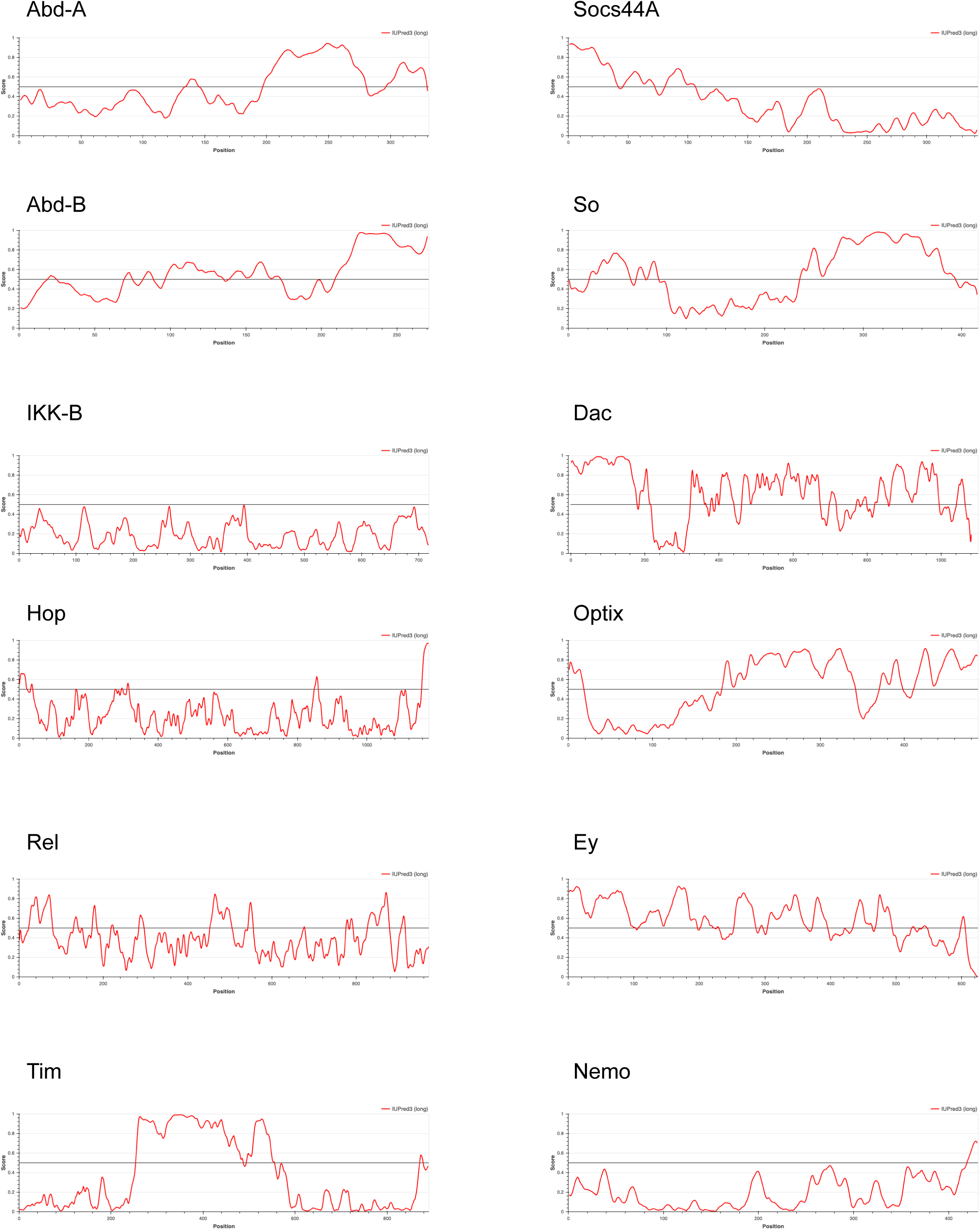
IUPRED3 prediction of disorder in the Abd-A, Abd-B, IKK-B, Hop, Rel, Tim, Socs44A, So, Dac, Optix, Ey, and Nemo.

